# Defective base excision repair in the response to DNA damaging agents in triple negative breast cancer

**DOI:** 10.1101/685271

**Authors:** Kevin J. Lee, Cortt G. Piett, Joel F Andrews, Elise Mann, Zachary D. Nagel, Natalie R. Gassman

## Abstract

DNA repair defects have been increasingly focused on as therapeutic targets. In hormone positive breast cancer, XRCC1-deficient tumors have been identified and proposed as targets for combination therapies that damage DNA and inhibit DNA repair pathways. XRCC1 is a scaffold protein that functions in base excision repair (BER) by mediating essential interactions between DNA glycosylases, AP endonuclease, poly(ADP-ribose) polymerase 1, DNA polymerase β (POL β), and DNA ligases. Loss of XRCC1 confers BER defects and hypersensitivity to DNA damaging agents. BER defects have not been evaluated in triple negative breast cancer (TNBC), for which new therapeutic targets and therapies are needed. To evaluate the potential of XRCC1 as an indicator of BER defects in TNBC, we examined XRCC1 expression and localization in the TCGA database and in TNBC cell lines. High XRCC1 expression was observed for TNBC tumors in the TCGA database and expression of XRCC1 varied between TNBC cell lines. We also observed changes in XRCC1 subcellular localization in TNBCs that alter the ability to repair base lesions and single-strand breaks. Subcellular localization changes were also observed for POL β that did not correlate with XRCC1 localization. Basal levels of DNA damage were also measured in the TNBC cell lines, and damage levels correlated with observed changes in XRCC1 expression, localization, and repair functions. The results confirmed that XRCC1 expression changes may indicate DNA repair capacity changes but emphasize that basal DNA damage levels along with expression and localization are better indicators of DNA repair defects. Given the observed over-expression of XRCC1 in TNBC preclinical models and the TCGA database, XRCC1 expression levels should be considered when evaluating treatment responses of TNBC preclinical model cells.

## Introduction

Defects in DNA damage response and repair are driving factors in carcinogenesis and key determinants in the response to chemotherapy. Breast cancers may display defects in DNA repair such as mutations in key DNA damage response and repair proteins such as breast cancer-susceptibility (*BRCA1/2*) and tumor suppressor protein p53 (*TP53*), and altered expression levels of DNA repair proteins thymine-DNA glycosylase (TDG) and poly(ADP-ribose) polymerase 1 (PARP1) [1–3]. Therapeutic outcomes may be improved by exploiting DNA repair defects present in cancer cells but absent in normal cells, as in the use of PARP-inhibitors (PARPi) in cancers that have BRCA1/2 deficiencies. However, characterization of DNA repair pathways often is lacking in preclinical models and cell lines. Examining DNA repair defects in preclinical models and patients is essential for evaluating the efficacy of therapeutic agents.

In breast cancer, DNA repair defects often extend beyond homologous recombination defects and observed changes in the expression of base excision repair (BER) proteins such as TDG and DNA polymerase beta (POL β). These defects imply that BER contributes to genomic instability and may alter therapeutic response [2, 3]. Single nucleotide polymorphisms in BER genes such as POL β, PARP1, and X-ray cross complementing protein 1 (XRCC1) may be linked to increased risk of developing breast cancer [4–7]. XRCC1 facilitates critical protein-protein interactions at the site of DNA damage with DNA glycosylases, AP endonuclease 1 (APE1), PARP1, POL β, and DNA ligase III (LIG3) and functions in the overlapping of BER and single-strand break repair (SSBR) pathways [8, 9]. XRCC1 also participates in double-strand break (DSB) repair through its interaction with PARP1 in the error-prone alternative nonhomologous end joining (NHEJ) [10–12], and with DNA LIG3 in nucleotide excision repair (NER) [13].

XRCC1 is ubiquitously expressed in normal tissues, but low levels of XRCC1 causing impaired BER may occur in terminally differentiated muscle cells and neurons [14, 15]. Variations in XRCC1 expression levels have been observed in breast cancer patient samples [16–18], and breast cancers with low expression levels of XRCC1 have been proposed as targets for PARPi treatment [16–19]. Defects in BER and XRCC1 may sensitize breast cancer cells to PARPi, similar to homologous recombination and BRCA1/2 defects [16, 18, 20, 21]. However, the mechanism of sensitization of XRCC1-deficient cancer cells to PARPi therapy remains unclear.

Defects in BER including XRCC1 expression level changes have not been explored in triple negative breast cancer (TNBC). TNBC may be aggressive and unresponsive to current treatments that target estrogen receptor (ER), progesterone receptor (PR), and human epidermal growth factor receptor 2 (HER2). However, TNBCs have a high prevalence of DNA repair defects, though not always associated with mutations in the *BRCA1/2* genes [1, 16–18, 22].

It is important to understand DNA repair defects that may promote resistance to available treatment. Characterizing and targeting XRCC1-related DNA repair defects in TNBCs may provide new treatment strategies for this aggressive breast cancer subtype. We examined XRCC1 expression levels and DNA repair functions in four TNBC model cell lines that are used commonly for preclinical screening of therapeutics. These models have higher basal levels of DNA damage that correlate with XRCC1 levels or XRCC1 cytosolic localization, and affect their sensitivity to DNA damaging agents. These observations lead us to hypothesize that XRCC1 expression and localization may be potential markers to develop treatment options for TNBC.

## Materials and Methods

### Cell culture

MDA-MB-157 (MDA-157), MDA-MB-231(MDA-231) and MDA-MB-468 (MDA-468), MCF10A, and HCC1806 cells were purchased from the American Type Culture Collection (ATCC #’s HTB-24, HTB-26, HTB-132, CRL-10317, and CRL-2335, respectively) within the last 12 months and passaged < 15 times for all experiments. Cells were tested biweekly during experiments using a mycoplasma detection kit to confirm absence of mycoplasma contamination using the Lonza MycoAlert^®^ (Lonza #LT07-318). HCC1806 cells were grown in RPMI 1640 Medium (Life Technologies #11875093) and supplemented with 10% Fetal Bovine Serum (FBS, Atlantic Biologicals Premium Select). MDA-157, MDA-231, MDA-468, and MCF10A cells were grown in DMEM High Glucose + GlutaMAX™ (Life Technologies #10566016) and supplemented with 1% sodium pyruvate (Life Technologies #11360070) and 10% FBS. Cells were maintained in a humidified 37°C incubator with 5% carbon dioxide.

### Fluorescence multiplex host cell reactivation

Fluorescence multiplex host cell reactivation (FM-HCR) assays were performed as described previously [23, 24]. Cells were seeded 48 hours before transfection into T25 tissue culture flasks (Thermofisher,156367) and collected for transfection at 85% confluence. Cells were electroporated using the Neon transfection system (ThermoFisher, MPK5000) at 1200 V, and 20 ms, with 2 pulses. Transfected cells were seeded into 12-well culture plates and harvested 24 hours after transfection for flow cytometric analysis. Reporter expression was reported as a percentage of fluorescent reporter protein expression from undamaged control plasmids in a second transfection that was also normalized for transfection efficiency. Fluorescent signal generated from the repair of the damaged and undamaged substrates was quantitated in each triplicate, and percent reporter expression was calculated as described previously [23, 24].

### Cytotoxicity

Cytotoxicity was determined with cell growth inhibition assays. Cells were seeded at 3 x 10^4^ cells per well in 6-well dishes. HCC1806 cells were treated the next day, and MDA-157, MDA-231, and MDA-468 were treated after 48 hours. All treatments were performed in triplicate with a minimum of 3 replicates each. Cells were treated with methyl methanesulfonate (MMS) (Sigma-Aldrich #129925) or potassium bromate (KBrO3) diluted in growth medium for 1 hour. Cells were washed in phosphate buffered saline (PBS, Cellgro #21-031-CV), growth medium was replaced, and the cells were incubated for 5 days. Cells were treated with olaparib (Selleck Chemicals #S1060) diluted in growth medium for 24 hours. Cells were washed with PBS, growth medium was replaced, and the cells were incubated for 5 days and counted with a Bio-Rad TC20 automated cell counter. Results were normalized to either vehicle control (olaparib) or untreated (MMS or KBrO_3_) cells and graphed to generate values of half maximal inhibitory concentration (IC_50_) values using GraphPad Prism software.

### Western blot

Western blot was performed as described previously [25]. Cells were grown in 150 mm dishes and cultured to 70-80% confluence. Cells were rinsed with PBS, scraped, stored overnight at −80°C, and lysed. Lysates were probed with antibodies diluted in 5% non-fat dry milk in Tris buffered saline (VWR #J640-4L) and 0.1% Tween20 (Fisher Scientific #BP337, TBS-T) and raised against PARP1 (Cell Signaling #9532) that was diluted 1:1000, XRCC1 (Fisher Scientific #MS434P1) that was diluted 1:500, p53 (Santa Cruz Biotechnology #sc-6243) that was diluted 1:1000, BRCA1 (Novus Biologicals #NBP1-45410) that was diluted 1:500, and glyceraldehyde 3-phosphate dehydrogenase (GAPDH) (Santa Cruz Biotechnology #sc-365062) that was diluted 1:1000.

### Immunofluorescence

Cells were plated in 8-well slides (Nunc Chambered Coverglass, ThermoFisher). Wells were coated with 250 μL EmbryoMax^®^ 0.1% Gelatin Solution (Millipore #ES-006-B) for 15-30 min at room temperature (RT ~ 23°C), EmbryoMax was removed and cells were added at 3 x 10^4^ cells per well and allowed to reach 70-80% confluence before fixation in 3.7% formaldehyde (Fisher Scientific #BP531) in PBS for 10 minutes at room temperature, then washed 3x in PBS. The cells were permeabilized using Permeabilization Buffer (Biotium #22016) for 10 minutes at room temperature, washed 3x in PBS, and blocked using 2% bovine serum albumin (BSA, Jackson ImmunoResearch #001-000-173) in PBS for 30 minutes at room temperature. Primary antibodies were diluted in 2% BSA in PBS and incubated for 1 hour at room temperature. XRCC1 (abcam #ab1838) was diluted 1:100; POL β (abcam #ab26343) was diluted 1:200; 53BP1 (Novus #NB100-304) was diluted 1:750; γH2AX-647 (Millipore #05-636-AF647) was diluted 1:750; Poly ADP-Ribose Polymer (PAR) (abcam #ab14460) was diluted 1:200; p53 (Life Technologies #MA514516) was diluted 1:200; and PARP1 (Santa Cruz Biotechnology #sc53643 clone C2-10) was diluted 1:200. After primary incubation, cells were washed 3x in PBS and secondary antibodies were diluted 1:2000 in 2% BSA and incubated for 45 min at room temperature in the dark. Secondary antibodies were either goat anti-Mouse Alexa Fluor 546 (Invitrogen #A11003), goat anti-Rabbit Alexa Fluor 546 (Invitrogen #A11010), goat anti-Mouse Alexa Fluor 488 (Invitrogen #A11001), or goat anti-Rabbit Alexa Fluor 488 (Invitrogen #A11008). One drop of NucBlue™ Fixed Cell Reagent (ThermoFisher #R37606) was added to each chamber and chambers were incubated for 15 min at room temperature. Cells were washed 2x in PBS and imaged immediately for analysis or stored at 4°C overnight. Fluorescence images were acquired using a Nikon A1r scanning confocal microscope with a Plan-Apochromat 20x/0.75 objective. For quantification, the region of interest (ROI) generator was used to automatically detect nuclei in the DAPI channel, and the mean intensity in the channel of interest was exported for analysis. For XRCC1 and POL β nucleus to cytoplasmic ratio (N/C), nuclei were defined automatically and the whole cell was drawn using Bezier function. The nucleus was then subtracted from the whole cell intensity to determine cytoplasmic intensity described previously [26, 27].

### 355 nm Laser microirradiation

Cells were plated as for immunofluorescence in 8-well chambered coverglass at 4 x 10^4^ cells per chamber. The next day, cells were treated with 10 μM bromodeoxyuridine BrdU (Sigma Aldrich #B5002) for 24 hours to sensitize the cells to microirradiation. Microirradiation was performed on a Nikon A1r scanning confocal microscope that was modified to include a 355 nm laser. Experiments were performed using an ultraviolet passing S Fluor 20x/0.75 objective as described previously [28, 29]. Samples were fixed with 3.7% formaldehyde at various time points and processed by immunofluorescence for XRCC1 fluorescence intensity. Intensity levels of damage foci were normalized by subtracting an adjacent region within the nucleus and reported as fluorescence intensity in arbitrary units (a.u.) as described [28].

### DNA Damage Analysis Utilizing RADD

Cells were plated in 8-well chambered coverglass, fixed, and permeabilized using the same procedure as for immunofluorescence. The Repair Assisted Damage Detection (RADD) assay was performed as described previously with minor modifications [30]. After permeabilization, cells were incubated with uracil DNA glycosylase (UDG) to remove uracil (NEB #M0304S), formamidopyrimidine [Fapy]-DNA glycosylase (Fapy-DNA glycosylase NEB #M0240S) to remove Fapy lesions, T4 Pyrimidine dimer glycosylase (T4PDG NEB #M-308S) to remove pyrimidine dimer lesions, endonuclease IV (Endo IV NEB #M0304S) to process oxidative damage, AP sites and modifies 3’ phosphates to 3’ OH, and endonuclease VIII (Endo VIII NEB #M0299S) to remove damaged pyrimidines diluted in 1X Thermpol buffer and incubated at 37 degrees C for 1 hour. This leaves gaps to be filled by DNA polymerase I Klenow large fragment (lacking 5’ to 3’ exonuclease activity) incubated with Digoxigenin-11-dUTP, alkali-labile (Dig) (Sigma-Aldrich #DIUTP-RO) at 37 degrees C for 1 hour. This allows the Dig-dUTP to be incorporated into the DNA. Cells were then washed in PBS, blocked using 2% BSA in PBS and Dig was then detected using an anti-Dig antibody (abcam #ab420 clone 21H8) at a dilution of 1:250 in 2% BSA in PBS for 1 hour at room temperature. Samples were then counterstained with Alexa Fluor goat anti-mouse 546 at 1:400 in 2% BSA in PBS for 45 min at room temperature and nuclei were stained as for immunofluorescence. Image analysis was performed as described previously [30]. MDA-231 cells were used to set the 546 nm laser gain, and the other cells were imaged at that gain. The control experiment was performed with the complete RADD cocktail and Klenow but without the insertion of Dig-dUTP. Antibody staining in this sample was used to ensure specificity of RADD measurements. A minimum of 200 cells per experiment were analyzed. The mean nuclear fluorescence intensity was normalized to the MDA-231 cells and reported as ratio of change over MDA-231.

### The Cancer Genome Atlas analysis

The Cancer Genome Atlas (TCGA) analysis was performed with an internet resource (UALCAN: http://ualcan.path.uab.edu) as previously reported by Chandrashekar et al [31] and plotted as a Box and Whisker to show data distribution.

### Statistical Analysis

Quantification of fluorescent images was performed in the Nikon Elements software package with a minimum of at least 150-200 cells per experiment, with each experiment performed at least three times. IC_50_ values were generated utilizing the GraphPad Prism software with three replicates for each concentration, repeated a minimum of three times, and average IC_50_ values reported. One-way ANOVA and means were compared with Dunnett post hoc analysis. Means were reported ± standard error of the mean (SEM).

## Results

### XRCC1 expression in the TCGA

The TCGA Breast Invasive Carcinoma dataset was analyzed using the UALCAN data portal [31] for XRCC1 expression. Despite previous publications indicating a loss in XRCC1 to be an important indicator for TNBC [16], XRCC1 transcript expression levels were significantly higher in primary tumor samples than in normal tissues with a p value of 1.6 x 10^-12^ (Fig 1A). Further analysis of XRCC1 expression across breast cancer types shows increased expression of XRCC1 in Luminal (p < 1 x 10^-12^) and TNBC (p < 0.001) tumor types (Fig 1B), but not in HER2 positive tumors.

**Fig 1.**
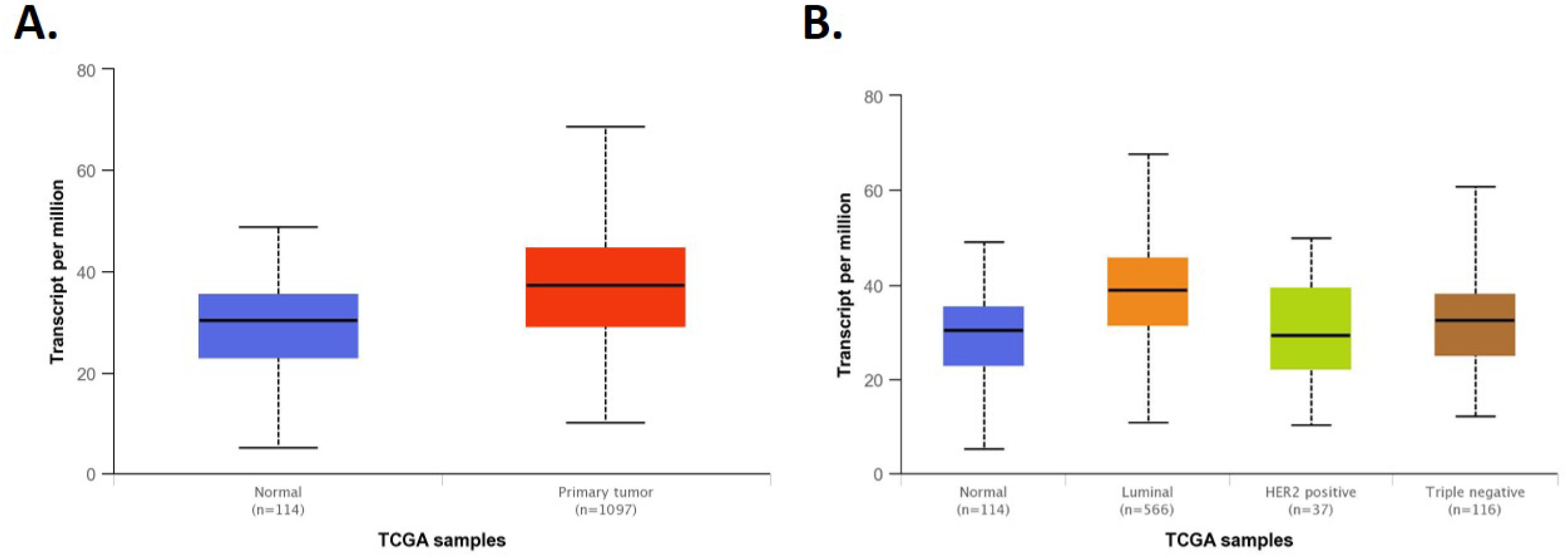
TCGA analysis of XRCC1. A) TCGA analysis using the UALCAN web interface [31] revealed XRCC1 transcript expression to be significantly higher in breast tumor tissue over normal tissue (p = 1.6 x 10^-12^). B) XRCC1 was increased in Luminal (p < 1 x 10^-12^) and TNBC (p < 0.001) tumor types compared to normal tissue using the same analysis interface.

### Base excision repair protein expression

To further examine XRCC1 expression and the expression of other key DNA repair proteins *in vitro* we utilized four highly characterized TNBC cell lines: MDA-157, MDA-231, HCC1806, and MDA-468 [32–34]. The expression levels of PARP1 were moderate and similar in MDA-157, HCC1806, and MDA-468 but lower in MDA-231 cells (Fig 2A). We observed low POL β protein levels in all cell lines with the highest level of POL β expression observed in MDA-468 cells, while MDA-231 and MDA-468 had similar high levels of p53 due to stabilizing mutations that also promote tumorigenesis through gain-of-function activities [35, 36], but we detected no p53 protein in MDA-157 cells, and low levels in HCC1806 cells that contained truncating mutations in p53 (Fig 2A, S1A Fig) [37–40]. Levels of XRCC1 expression were lowest in MDA-157 cells similar to non-tumorigenic epithelial MCF10A cells, higher in MDA-231 and HCC1806 cells, and the highest in MDA-468 cells (Fig 2B, quantified in S1A Fig). We observed a similar pattern of BRCA1 expression that correlated to XRCC1 expression in the absence of any reported BRCA1 mutation in these cell lines (S1B Fig) [33].

**Fig 2.**
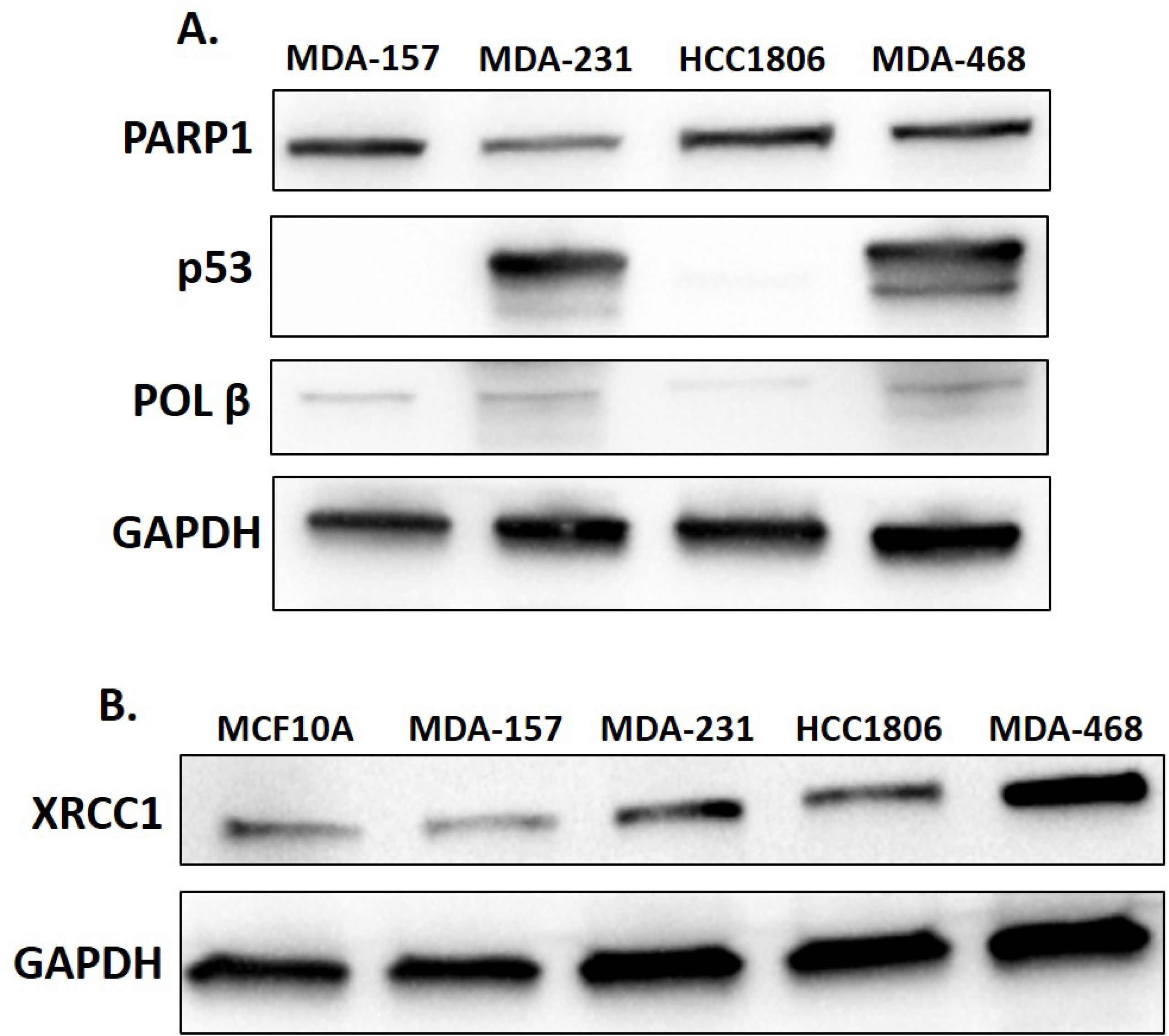
Base excision repair proteins in TNBC cell lines. A) Lysates of MDA-MB-157 (MDA-157), MDA-MB-231 (MDA-231), HCC1806, and MDA-MB-468 (MDA-468) were probed by western blot for the expression of PARP1, p53, POL β, and GAPDH as a loading control. B) Lysates of MCF10A, MDA-157, MDA-231, HCC1806, and MDA-468 were probed by western blot for the expression of XRCC1 with GAPDH serving as loading control.

### Subcellular localization of critical base excision repair proteins

For effective DNA repair and maintenance of genomic fidelity, the proteins involved must be localized in the nucleus. We examined the expression and nuclear localization of XRCC1 and POL β using immunofluorescence. Previous studies showed that POL β stability may depend on interactions with XRCC1 [41–43], and XRCC1 localization has been previously observed to be primarily nuclear [26]. POL β functions in both nuclear and mitochondrial BER and is observed within the cytoplasm, mitochondria, and nucleus of cells [27, 44].

We assessed the nuclear to cytoplasmic ratio (N/C) of XRCC1, with a value of 1 representing equal distribution between the nucleus and cytoplasm, while a value of less than one represents exclusion from the nucleus, and a value greater than one represents predominantly nuclear localization. The MDA-157 cells exhibited relatively equal distribution of XRCC1 between the nucleus and cytoplasm with a N/C ratio of 1.01 ± 0.03 (Fig 3A and 3C). MDA-231 showed higher nuclear content of XRCC1 with a N/C ratio of 2.02 ± 0.12, though some cytoplasmic content is still observed. HCC1806 cells had partial exclusion of XRCC1 from the nucleus with a N/C ratio of 0.79 ± 0.02 (Fig 3A and 3C). MDA-468 cells had the highest N/C ratio of the cell lines tested (2.36 ± 0.12) with XRCC1 almost exclusively in the nucleus (Fig 3A and 3C).

**Fig 3.**
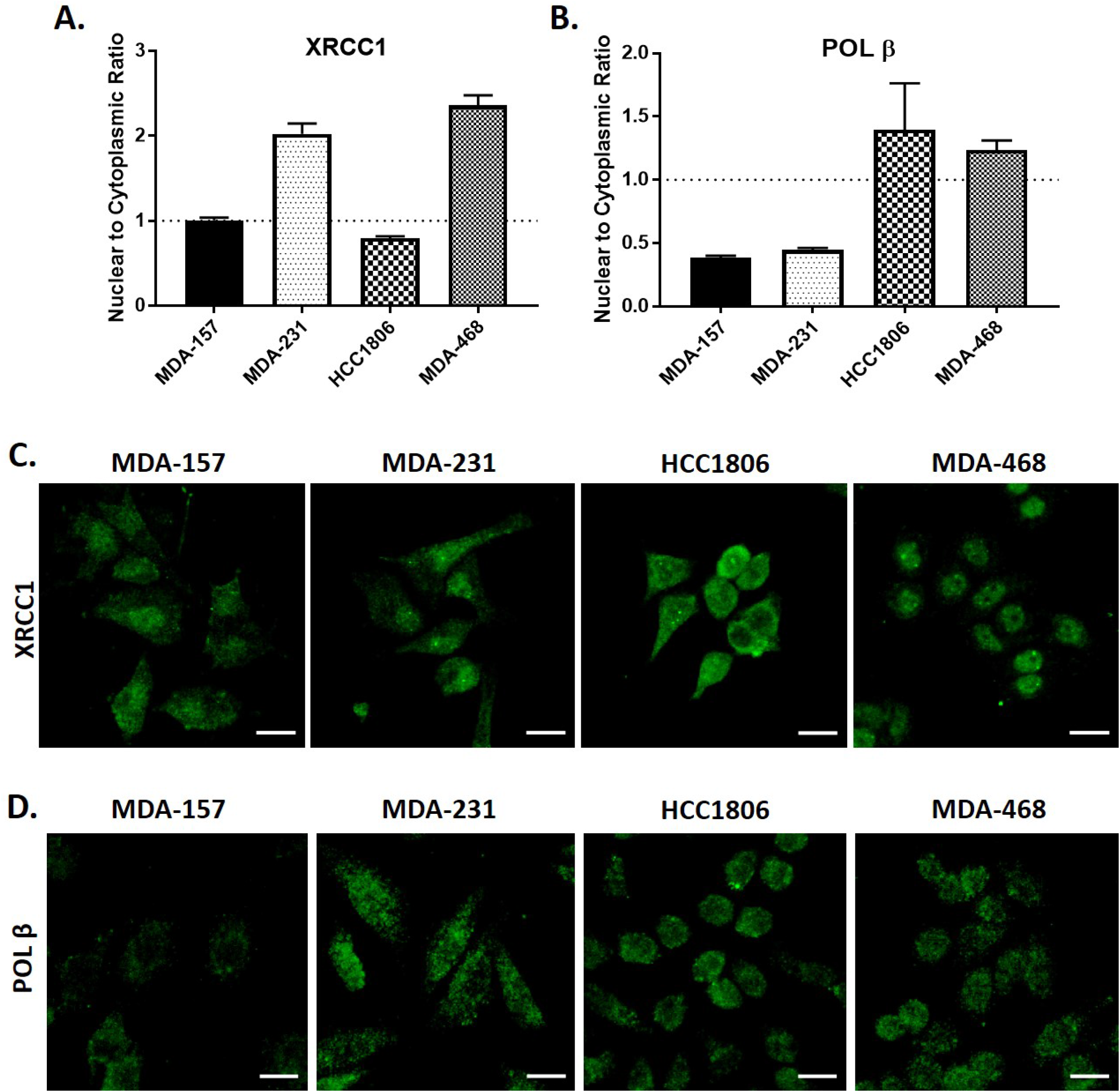
Subcellular localization and quantification of BER proteins. A) XRCC1 nuclear to cytoplasmic ratio (N/C) is reported as 1 representing equal distribution (MDA-157), > 1 representing nuclear localization (MDA-231 and MDA-468), and < 1 representing nuclear exclusion (HCC1806). B) N/C ratio of POL β. C) Representative images of localization of XRCC1 in TNBC cell lines. D) Representative images of localization of POL β in TNBC cell lines. Scale bar = 20 μm.

Given the diminished nuclear localization of XRCC1 in HCC1806 cells, we also examined the nuclear content of POL β across the cell lines. Consistent with western blot results, POL β staining was low across all cell lines, and there was variability in the nuclear content of POL β between cells (Fig 3B and 3D, S2 Fig). There was relatively lower nuclear content of POL β in MDA-157 and MDA-231 cells with N/C ratios of 0.39 ± 0.02 and 0.45 ± 0.02, respectively (Fig 3B), in contrast to the nuclear content of XRCC1 in these cell lines. HCC1806 showed nuclear enrichment of POL β with an N/C of 1.40 ± 0.37, despite XRCC1 being predominately cytoplasmic (Fig 3B and 3D). MDA-468 cells also had enriched nuclear POL β with an N/C of 1.24 ± 0.08.

Given the unexpected distribution of BER factors, we also examined cellular levels of PAR, which is polymerized by PARP1 at sites of DNA damage, and increases in the presence of BER defects [42, 45]. Nuclear levels of PAR were varied, with MDA-157 cells having the highest nuclear intensity of PAR and HCC1806 cells unexpectedly having the lowest nuclear intensity of PAR, despite XRCC1 being largely cytoplasmic (Fig 4A).

**Fig 4.**
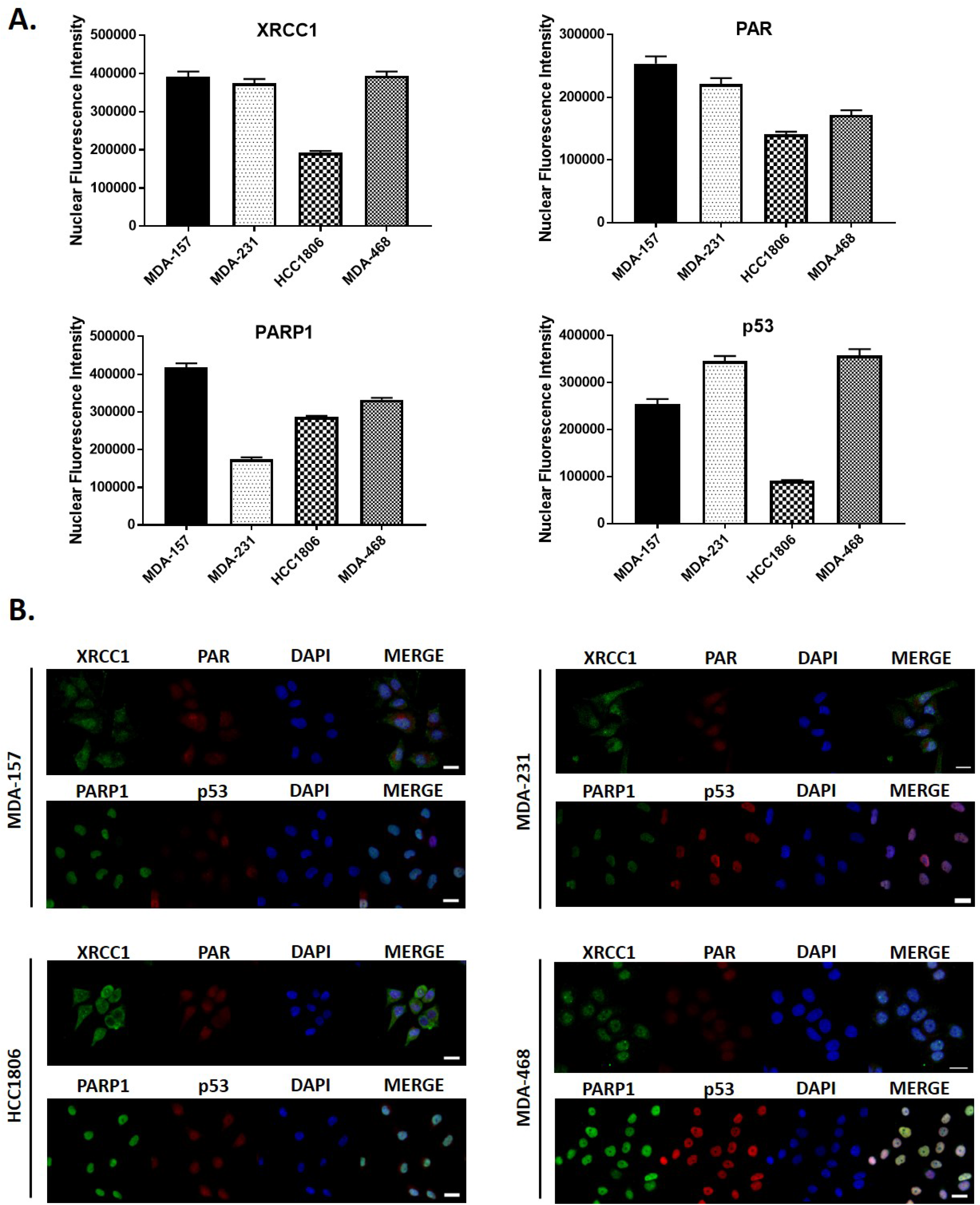
Nuclear intensity of key BER proteins. A) Quantification of mean nuclear fluorescence intensity is shown for XRCC1, PAR, PARP1, and p53. B) Representative fluorescent images used for quantification in A for XRCC1, PAR, PARP1, and p53 for all cell lines tested. Scale bar = 20 μm.

We then examined the nuclear content of PARP1 and p53. MDA-157 contained roughly 2.4-fold more nuclear PARP1 than MDA-231 cells. HCC1806 had 1.6-fold and MDA-468 1.9-fold more nuclear PARP1 than MDA-231 cells (Fig 4A). PARP1 was highly localized to the nucleus in all cell lines (Fig 4B). We observed that p53 nuclear localization was consistent with mutational status, with HCC1806 and MDA-157 displaying weak nuclear staining, in contrast with strong nuclear staining of p53 in MDA-231 and MDA-468 cells, consistent with Western blot data (Fig 2). Representative images of all cell lines are shown in Fig 4B, with XRCC1 and PARP1 shown in green, while PAR and p53 are shown in red.

### Basal levels of DNA damage in TNBC cell lines

Based on the altered protein expression and localization of BER proteins in TNBC cells, we evaluated cells for variation in levels of basal DNA damage using the broad spectrum DNA damage assay, Repair Assisted Damage Detection (RADD) [30]. RADD uses bacterial DNA glycosylases to detect and excise abasic sites, oxidative base lesions, bulky DNA adducts, and thymine dimers (see Material and Methods). Then the gaps in the DNA where the lesions were excised and any strand breaks occur are then tagged by Klenow polymerase with a Digoxigenin-11-dUTP. Damage sites are detected by immunofluorescence as described in the materials and methods [30]. RADD was applied to the TNBC cell panel, and the nuclear fluorescence intensity of basal DNA damage quantified (Fig 5, S3 Fig). MDA-157 and MDA-231 cells had similar low levels of basal DNA damage (Fig 5A). However, HCC1806 cells had 1.4-fold more DNA damage over MDA-231 (P = 0.0501) and MDA-468 cells had 1.8-fold more damage than MDA-231 cells (P < 0.01) (Fig 5A). Representative images of nuclear fluorescence of all cell lines is shown in Fig 5B.

**Fig 5.**
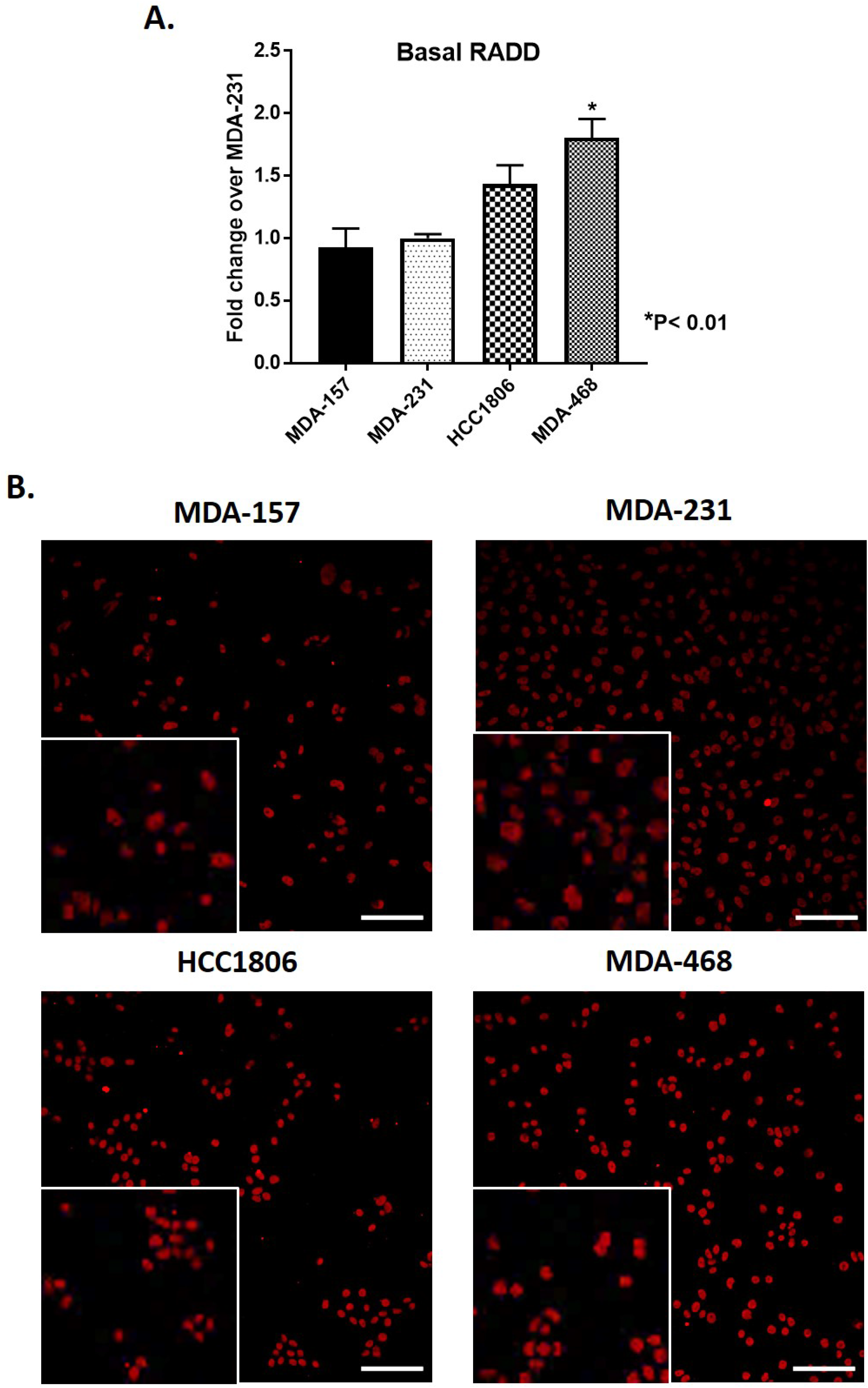
Basal DNA damage in TNBC cell lines. A) Quantification of images in B normalized to MDA-231 show an increase in DNA damage in HCC1806, and a significantly higher amount (P < 0.01) of DNA damage for MDA-468. B) Representative images of basal levels of DNA damage as measure by RADD in MDA-157, MDA-231, HCC1806, and MDA-468 with insets showing blown up cropped images. Scale bar = 100 μm.

### XRCC1 recruitment to single-strand breaks is altered in TNBC cells

The varied levels of basal DNA damage and altered expression and localization of XRCC1 indicated that DNA repair functions of XRCC1 may be altered in TNBC cells. As XRCC1 has no known enzymatic activity, we assessed DNA repair activity by examining the recruitment and retention of XRCC1 at sites of DNA damage induced by laser microirradiation. Single-strand breaks were induced in a subnuclear region by microirradiation with a 355 nm laser, and we assessed XRCC1 recruitment and retention to the damage sites at various time points after inducing damage [28, 29].

As shown in Fig 6, XRCC1 recruitment over time is graphed on the left, with corresponding images of peak XRCC1 recruitment foci shown on the right. XRCC1 was recruited to the DNA damage spot in MDA-157 cells with an average intensity of 707 a.u. that was resolved within 5 minutes, with 50% retention at 1.7 minutes (Fig 6A). In contrast, MDA-231 cells had two peaks of XRCC1 recruitment at 364 a.u. within 0.5 minutes that resolved by 3 min, and at 8 minutes (Fig 6B). XRCC1 in HCC1806 cells had weak recruitment to the damage spot, peaking only at 258 a.u., likely due to mislocalization of XRCC1 as indicated in Fig 3A (Fig 6C). In MDA-468 cells, XRCC1 recruitment was rapid and robust, peaking at 1025 a.u. within 0.5 minutes and 50% retention at 2.2 minutes (Fig 6D), and XRCC1 was rapidly dissociated from the damage site, with complete resolution of the damage foci within 3 minutes of microirradiation (Fig 6D). Double-strand breaks (DSBs) were absent in all cell lines as confirmed by the lack of co-localization of DSB markers 53BP1 and γH2AX at 10 minutes after microirradiation, except for a slight increase of γH2AX at 10 minutes in HCC1806 cells (S4 Fig, and S5 Fig).

**Fig 6.**
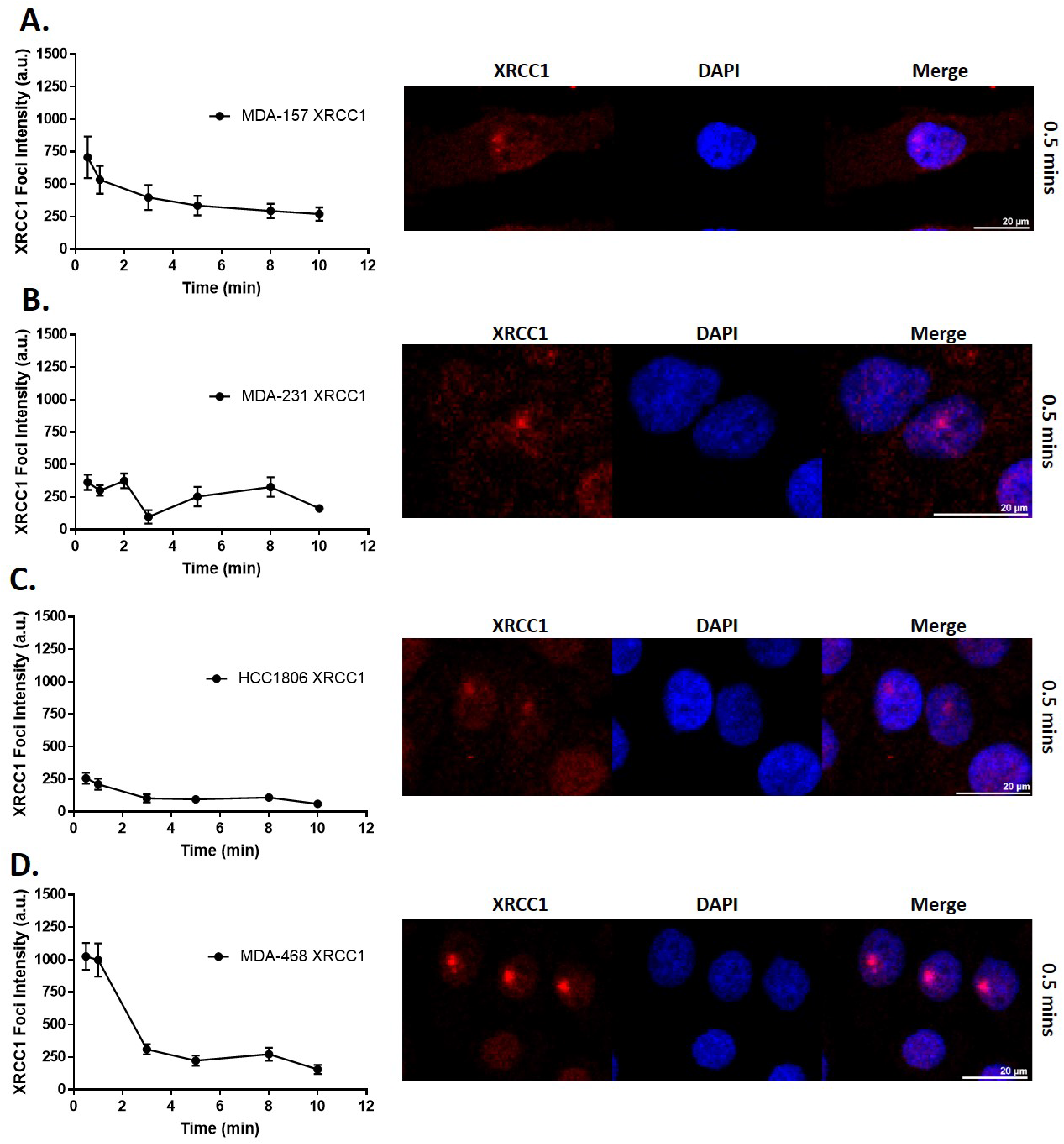
XRCC1 recruitment following 355 nm laser microirradiation in TNBC cell lines. Microirradiation with a 355 nm laser occurred and cells were fixed at the indicated time points and processed for immunofluorescence of XRCC1. Quantification over time is shown on the left with a representative image from 0.5 min shown on the right for A) MDA-157, B) MDA-231, C) HCC1806, and D) MDA-468. A minimum of 18 cells were irradiated over three separate experiments and quantified as described in the materials and methods section and reported as XRCC1 Foci Intensity in arbitrary units (a.u.). Scale bar = 20 μm

### Base excision repair capacity is altered in TNBC cells

Given the BER defects observed through expression, localization, and response to induced DNA damage, we utilized a flow cytometric host cell reactivation assay (FM-HCR) to assess the repair capacities of TNBC cells across BER substrates [23, 24]. The FM-HCR assays measured DNA repair capacity using transiently transfected fluorescent reporter plasmids. By reporting repair of each DNA lesion in a different fluorescent channel, simultaneous measurements were made for multiple DNA repair pathways to assess the diverse BER capacity in TNBC cells. The BER reporters include GFP_hypoxanthine:T, which primarily reports AAG (also known as MPG) glycosylase-mediated excision of hypoxanthine opposite thymine (Fig 7A); mPlum_A:8-oxo-dG, which reports MUTYH glycosylase catalyzed excision of adenine opposite 8-oxo-2’-deoxyguanosine (8-oxo-dG) (Fig 7B); mOrange_8-oxo-dG:C, which reports the activity of several glycosylases (OGG1, NEIL1, NEIL2) that excise 8-oxo-dG opposite cytosine (Fig 7C); and BFP_Uracil:G, which primarily reports UNG glycosylase catalyzed excision of uracil opposite guanine (Fig 7D). mPlum_O^6^-methylguanine:C was used to assess removal of O^6^-methylguanine opposite thymine by MGMT (Fig 7E).

**Fig 7.**
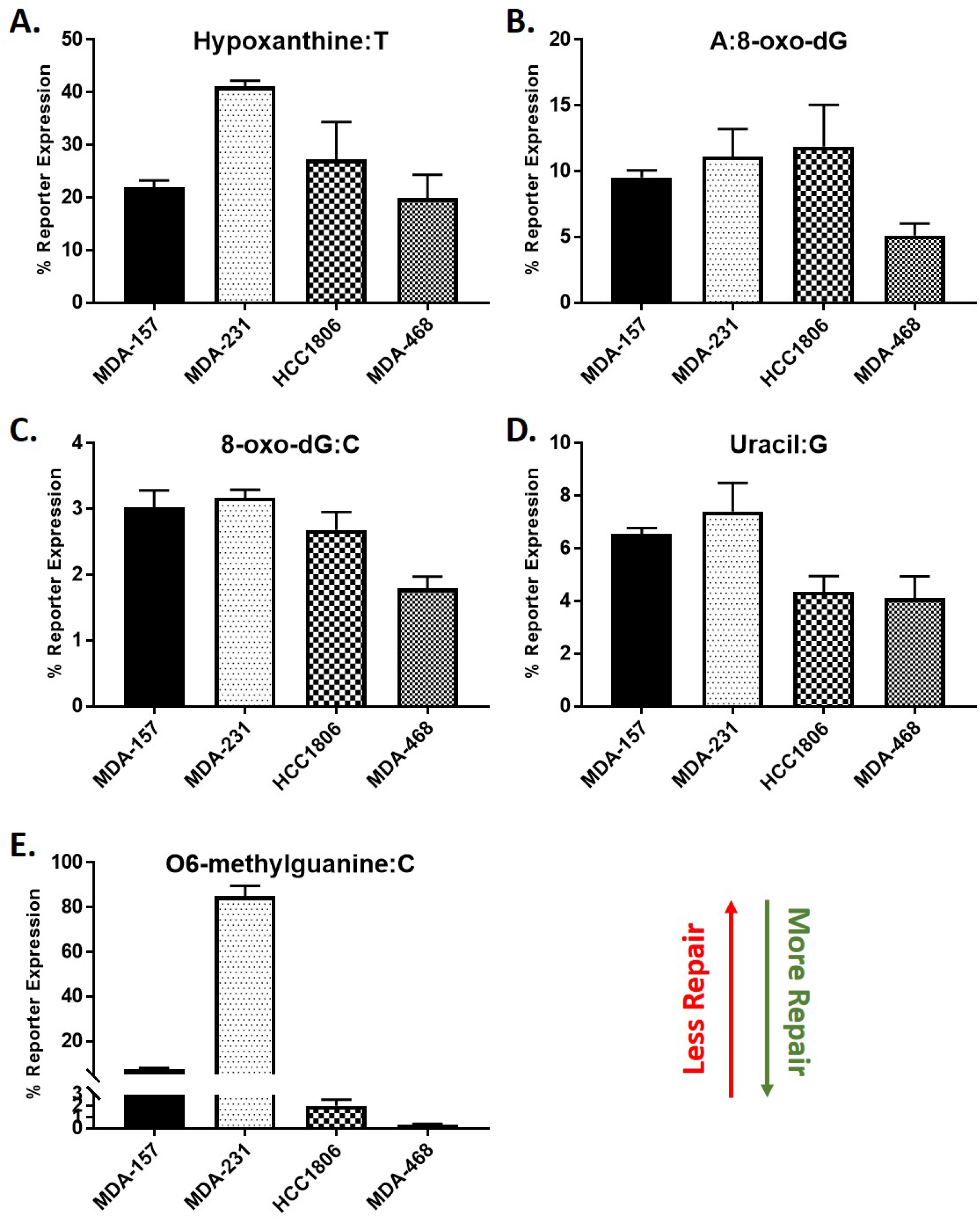
FM-HCR analysis of DNA repair capacity in TNBC cell lines. Cells were transfected with fluorescent reporter plasmids containing the indicated DNA lesions, A) Hypoxanthine:T, B) A:8-oxo-dG, C) 8-oxo-dG:C, D) Uracil:G, E) O6-methylguanine:C, as well as an undamaged plasmid to normalize for transfection efficiency. DNA repair capacity is inversely proportional to % reporter expression.

In the selected TNBC cells, MDA-468 cells were the most BER proficient with efficient repair observed for all BER substrates. MDA-468 also efficiently repaired O^6^-methylguanine by direct reversal. MDA-231 cells were defective for repair of O^6^-methylguanine, and overall least proficient at repairing BER substrates. MDA-157 cells repaired most BER substrates except for 8-oxo-dG:C and uracil, and showed intermediate repair efficiency for O^6^-methylguanine, repair of hypoxanthine:T with high efficiency similar to MDA-468, and repair of Uracil:G, 8-oxo-dG:C and A:8-oxo-dG with low efficiency similar to MDA-231 cells. HCC1806 cells repaired O^6^-methylguanine efficiently, and had low repair capacity for all BER substrates except uracil, for which repair proficiency matched MDA-468 cells.

### DNA damaging agents induce selective cytotoxicity in TNBC cells

We then analyzed sensitivity of the four TNBC cell lines to clinical DNA damaging agents. Toxic effects of cisplatin and doxorubicin have been widely reported for these cell lines with a large range of IC_50_ values observed (Table 1). BER proteins, including XRCC1, are involved in the repair of the platinum-DNA adducts, strand breaks, and DNA-protein crosslinks induced by these agents, though other repair pathways are more dominant [46–49].

**Table 1.**
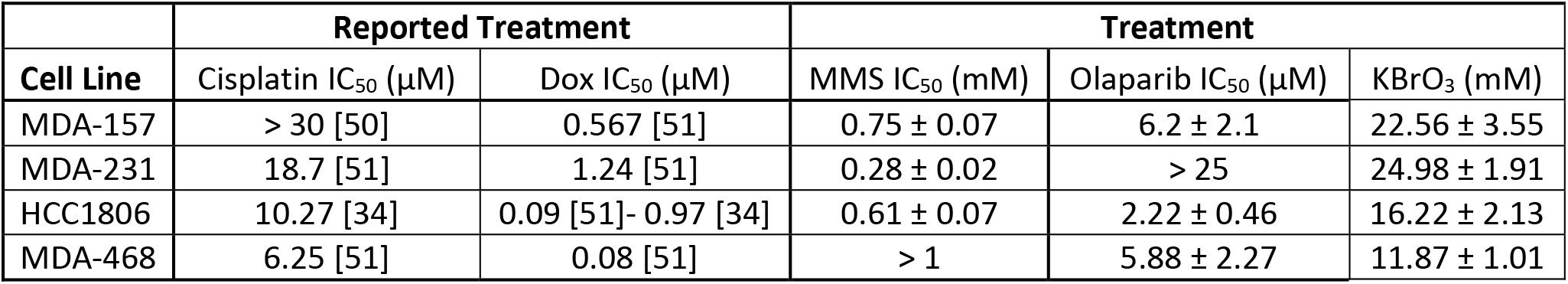
Sensitivity of TNBC cell lines to DNA damaging agents. Cisplatin and doxorubicin (Dox) IC_50_’s previously reported.

| Cell Line | Reported Treatment |  | Treatment |  |  |
| --- | --- | --- | --- | --- | --- |
|  | Cisplatin IC<sub>50</sub> (μM) | Dox IC<sub>50</sub> (μM) | MMS IC<sub>50</sub> (mM) | Olaparib IC<sub>50</sub> (μM) | KBrO<sub>3</sub> (mM) |
| MDA-157 | > 30 [50] | 0.567 [51] | 0.75 ± 0.07 | 6.2 ± 2.1 | 22.56 ± 3.55 |
| MDA-231 | 18.7 [51] | 1.24 [51] | 0.28 ± 0.02 | > 25 | 24.98 ± 1.91 |
| HCC1806 | 10.27 [34] | 0.09 [51]- 0.97 [34] | 0.61 ± 0.07 | 2.22 ± 0.46 | 16.22 ± 2.13 |
| MDA-468 | 6.25 [51] | 0.08 [51] | > 1 | 5.88 ± 2.27 | 11.87 ± 1.01 |

We determined TNBC cell line sensitivity to the DNA methylating agent methyl methanesulfonate (MMS). BER is responsible for the resolution of alkylation base damage induced by MMS, and XRCC1-deficient cells are hypersensitive to MMS [52]. We determined the MMS sensitivity (IC_50_) of the four TNBC cell lines. MDA-468 cells were the most resistant with IC_50_ greater than 1 mM MMS, followed by MDA-157 (0.75 mM) and HCC1806 cells (0.61 mM), MDA-231 cells were the most sensitive to MMS with an IC_50_ of 0.28 mM (Table 1).

Sensitivity to the PARPi olaparib was examined. PARP1 recognizes abasic sites and single-strand breaks and synthesizes poly(ADP-ribose) (PAR) polymers to recruit additional BER proteins such as XRCC1, POL β, and DNA LIG3 for repair [8, 21]. Olaparib inhibits PAR production thereby reducing BER mediating repair [48, 53].

Cells were treated with increasing concentrations of olaparib up to 25 μM. MDA-231 cells were most resistant to olaparib treatment up to the highest concentration tested, 25 μM (Table 1). MDA-157 and MDA-468 cells were sensitive to olaparib treatment with IC_50_ values of 6.2 and 5.88 μM, respectively, while HCC1806 cells were the most sensitive to olaparib treatment with an IC_50_ of 2.22 μM (Table 1).

BER is also responsible for the repair of oxidative DNA lesions [54, 55]. TNBC cell sensitivity to the oxidizing agent potassium bromate (KBrO_3_ [54]) was tested at increasing concentrations up to 25 mM. MDA-231 and MDA-157 exhibited IC_50_ values of 24.98 mM and 22.56 mM, respectively, with no significant difference between the two. HCC1806 cells were sensitive to KBrO_3_ with an IC_50_ value of 16.22 mM representing a significant difference compared to MDA-231 (P < 0.05) (Table 1). MDA-468 cells were also sensitive to KBrO_3_ with an IC_50_ of 11.87 mM representing a significant difference compared to MDA-231 cells (P< 0.01) (Table 1).

## Discussion

Characterization of DNA repair defects has been increasingly clinically focused with gene expression panels and biomarkers sought to predict patient response and improve therapeutic outcomes. However, characterization of DNA repair pathways is often lacking in preclinical models and cell lines. Examining DNA repair defects in both preclinical models and patients is essential for evaluating the efficacy of therapeutic agents. Several studies have noted deficiency in XRCC1 in breast cancers and proposed deficiency in XRCC1 and defects in BER as targets for therapeutic intervention [16, 18, 19]. However, characterization of BER defects, particularly those linked to XRCC1 in breast cancer has been lacking. Given the aggressiveness of TNBC and the poor treatment options for recurrent TNBC, identifying and characterizing BER defects which may promote resistance to standard of care treatment is critical. Characterization of these defects has the potential to guide treatment options to increase survival.

While expression changes in PARP1 and POL β have been noted previously in TNBC, these expression changes have failed to serve as effective biomarkers for therapeutic response [1–3]. So we evaluated XRCC1 expression in the TCGA Breast Invasive Carcinoma dataset using the UALCAN data portal [31]. Unlike previous reports, over-expression of XRCC1 was observed in this dataset similar to the over-expression observed in the four preclinical models used in this study. XRCC1 transcript expression levels were higher in tumor samples than in normal tissues, and in Luminal and TNBC subtypes compared with normal tissue. Pathology images from the TCGA dataset also showed high protein content in breast carcinomas consistent with the high transcript levels [56]. Therefore, XRCC1 over-expression may be a more relevant to examining DNA repair defects in TNBC than XRCC1 deficiency.

To explore this hypothesis, we examined XRCC1 in four TNBC preclinical models to determine if XRCC1 expression changes account for varied responses to DNA damaging agents frequently observed [32–34]. XRCC1 protein levels varied across the four common preclinical model cell lines, and more importantly, XRCC1 localization also varied among the cell lines. XRCC1 functions as a scaffold protein in BER is well described and its localized into the nuclear compartment [26, 57, 58]. There is no evidence that XRCC1 is involved in mitochondrial BER, and mislocalization of XRCC1 into the cytoplasm by the single nucleotide polymorphism R280H has been associated with defective nuclear BER [59–61], though all cell lines used in this study lack this particular single nucleotide polymorphism (data not shown, COSMIC database). We observed significant cytoplasmic content of XRCC1 except in MDA-468 cells, in which XRCC1 was almost exclusively located in the nucleus (Fig 3A and 3C). The low nuclear content of XRCC1 in HCC1806 cells contributed to the sensitivity of this cell line to DNA damaging agents (Table 1). This loss in repair capacity is also supported by the low recruitment and retention of XRCC1 to laser-induced DNA damage and low repair of all BER substrates except uracil in the FM-HCR assay.

Since XRCC1 stabilizes the expression of POL β [41–43], we examined POL β expression between the cells lines. POL β also varied between the cell lines, with higher XRCC1 levels correlating with higher POL β protein content. However, the TNBC cells also showed altered localization of POL β that did not correlate with altered XRCC1 localization. The MDA-157 and MDA-231 cells showed low nuclear content of POL β, whereas HCC1806 and MDA-468 showed nuclear POL β enrichment [27, 44]. Loss of POL β does not induce the same level of hypersensitivity to DNA damaging agents as loss of XRCC1 [52, 62, 63]. However, loss of nuclear POL β may reduce efficiency of repair and likely contributes to reduced BER capacity and sensitivity of MDA-157 cells to MMS and other DNA damaging agents (Fig 7, Table 1).

The consequences of DNA repair defects in TNBC cells were observed with a broad spectrum method of detecting DNA damage [30]. Levels of XRCC1 (Fig 2) were correlated with increased levels of basal DNA damage. Low levels of XRCC1 in MDA-157 and MDA-231 cells corresponded to low levels of basal DNA damage, similar to that in the non-tumorigenic epithelial cell line MCF10A (S3 Fig). However, HCC1806 and MDA-468 cells had increased XRCC1 levels that corresponded to higher levels of DNA damage (Fig 5). We expected high levels of DNA damage in HCC1806 cells where XRCC1 is mislocalized to the cytoplasm. However, the high levels of DNA damage in MDA-468 cells, and the lack of known *BRCA1* mutations and dependence on mutant p53 for survival [36] may indicate a previously unidentified DNA repair defect in these cells or elevated levels of on-going repair that may cause increased expression of XRCC1, PARP1, and POL β.

Elevated levels of BER in MDA-468 cells are consistent with the proficiency of these cells to repair BER substrates (Fig 7) by the rapid and robust recruitment of XRCC1 to laser induced DNA damage (Fig 6), and the resistance of these cells to MMS (Table 1). The recruitment of XRCC1 was greater in MDA-468 than MDA-157 and HCC1806 cells, in which defects in XRCC1 expression and localization were observed. MDA-468 cells also recruited XRCC1 more robustly than MDA-231 cells, which had two recruitment waves of XRCC1 similar to a recent report examining XRCC1 in U2OS cells [64]. Interestingly, the second wave of XRCC1 recruitment observed in the U2OS cells was observed after CRISPR editing to make the cell XRCC1 deficient. It is unclear if second wave of recruitment is due to inefficient recruitment of POL β to the damage site from the low nuclear content of these cells (Fig 6) [42], or could possibly be due to a more complex BER pathway that is needed for correct end-tailoring of nicked DNA before gap-filling by the polymerase [28]. Further work is needed to better understand these recruitment dynamics.

The inefficiency in BER repair by MDA-231 cells was confirmed by the FM-HCR assays (Fig 7) and high sensitivity of MDA-231 cells to MMS (Table 1). The MDA-231 cells were not sensitive to olaparib alone and the least sensitive to KBrO_3_ of the lines tested, despite the defect in BER (Table 1). The sensitivity to olaparib may be reduced due to the lower levels of its target PARP1 observed in MDA-231 cells (Fig 2), while the lack of sensitivity to KBrO_3_ is likely due to the fact that while the MDA-231 cells are the least efficient in BER XRCC1 is still able to recruit to sites of damage (Fig 6). As the levels of DNA damage observed in this cell line are more comparable to BER competent MDA-157, low levels of PARP1 and the gain-of-function mutation in p53 observed in MDA-231 cells likely promote alternative repair pathways to reduce basal damage levels (Fig 5) [33]. Competition or ineffective signaling by these alternative repair pathways also may explain the altered recruitment profile for XRCC1 in the MDA-231 cells.

Although MDA-468 and MDA-231 cells represented the extremes of BER competencies, the observed sensitivity of MDA-157 and HCC1806 cells to MMS likely resulted from the observed defects in localization of BER proteins, which has not been previously assessed (Fig 3).

Characterization of XRCC1 expression in these preclinical models demonstrates that protein expression, localization, and the basal DNA damage levels are important indicators of DNA repair capacities. Despite the high proficiency for BER observed by FM-HCR, MDA-468 cells are well documented to be sensitive to DNA damaging agents [32, 51, 65]. Over-repair by the BER pathway and high basal DNA damage levels in these cells may generate too many cytotoxic intermediates that promote cell death and increase cellular sensitivity to DNA damaging agents, such as KBrO_3_ (Table 1), rendering these cells ineffective model for drug evaluation. The present data also demonstrate that defects in BER are prominent in the commonly used TNBC preclinical models. These BER defects have not been characterized or examined previously and need to be investigated in patient tumor samples.

In summary, XRCC1 expression and localization may vary and serve as critical indicators of BER defects in TNBCs and likely other cancers, however, XRCC1 in addition to other factors should be considered when analyzing BER capacity or defects in TNBC and should be analyzed in the context of the expression and localization of other BER proteins, such as POL β. Further, our work demonstrates the importance of assessing basal levels of DNA damage and DNA repair protein localization in cells and tissues may better predict treatment outcomes when DNA repair defects are present.

## Acknowledgements

This work was supported by R21ES028015 from National Institutes of Health, National Institute of Environmental Health Sciences and ONR Award N00014-16-1-3041. The ZDN lab is supported by SBDR P01 CA09584, from National Institutes of Health. The authors would like to thank Dr. Robert van Waardenburg for his review and comment on this work.

## Contributions

KJL and NRG designed the project and performed confocal microscopy experiments. KJL performed cytotoxicity assay and microirradiation assays. EM and NRG performed nuclear localization studies. JFA developed analysis software for microirradiation experiments. CGP and ZDN performed FM-HCR experiments and analysis. KJL prepared the figures and tables. KJL, NRG, CGP, and ZDN wrote the manuscript. All authors reviewed and approved the manuscript.

## Supplemental Figure Captions

S1 Fig. A) Western blots from Fig 1A, Fig 1B, and S1B Fig were quantified, normalized to loading control, and reported as fold change relative to MDA-231. ND = no signal detected. B) Lysates for MDA-157, MDA-231, HCC1806, and MDA-468 were probed for BRCA1, with GAPDH serving as a loading control.

S2 Fig. A) Mean nuclear fluorescence intensity of POL β. B) Representative images of POL β of all cell lines tested. Scale bar = 20 μm.

S3 Fig. Basal DNA damage in TNBC cell lines in comparison to MCF10A. A) Replotting data from Fig 4A in the presence of MCF10A shows similar levels of DNA damage to that of MDA-157 and MDA-231. B) Representative images of basal levels of DNA damage as measure by RADD including MCF10A. Scale bar = 100 μm.

S4 Fig. Double strand break markers post microirradiation. DSB markers 53BP1 (Green) and γ-H2AX (Violet) were stained by immunofluorescence at 10 min after micro-irradiation and representative images are shown for A) MDA-157, B) MDA-231, C) HCC1806, and D) MDA-468. Scale bar = 20 μm.

S5 Fig. Fluorescence intensity of double-strand break markers from S4 Fig. A) Foci Intensity for γ-H2AX (Left), and 53BP1 (Right) for MDA-157, MDA-231, HCC1806, and MDA-468. B) Mean ± SEM for γ-H2AX and 53BP1 from S5A Fig.

S6 Fig. FM-HCR analysis from Fig 6 including the non-tumorigenic cell line MCF10A.

## References

1. Li SX, Sjolund A, Harris L, Sweasy JB. DNA repair and personalized breast cancer therapy. Environ Mol Mutagen. 2010;51(8-9):897–908. doi: 10.1002/em.20606. PubMed PMID: 20872853; PubMed Central PMCID: PMCPMC2962983.

2. Wolf DM, Yau C, Sanil A, Glas A, Petricoin E, Wulfkuhle J, et al. DNA repair deficiency biomarkers and the 70-gene ultra-high risk signature as predictors of veliparib/carboplatin response in the I-SPY 2 breast cancer trial. NPJ Breast Cancer. 2017;3:31. doi: 10.1038/s41523-017-0025-7. PubMed PMID: 28948212; PubMed Central PMCID: PMCPMC5572474.

3. Carter SL, Eklund AC, Kohane IS, Harris LN, Szallasi Z. A signature of chromosomal instability inferred from gene expression profiles predicts clinical outcome in multiple human cancers. Nat Genet. 2006;38(9):1043–8. doi: 10.1038/ng1861. PubMed PMID: 16921376.

4. Karahalil B, Bohr VA, Wilson DM, 3rd. Impact of DNA polymorphisms in key DNA base excision repair proteins on cancer risk. Hum Exp Toxicol. 2012;31(10):981–1005. doi: 10.1177/0960327112444476. PubMed PMID: 23023028; PubMed Central PMCID: PMCPMC4586256.

5. Abdel-Fatah TM, Russell R, Agarwal D, Moseley P, Abayomi MA, Perry C, et al. DNA polymerase beta deficiency is linked to aggressive breast cancer: a comprehensive analysis of gene copy number, mRNA and protein expression in multiple cohorts. Mol Oncol. 2014;8(3):520–32. doi: 10.1016/j.molonc.2014.01.001. PubMed PMID: 24462520; PubMed Central PMCID: PMCPMC5528629.

6. Jiang J, Zhang X, Yang H, Wang W. Polymorphisms of DNA repair genes: ADPRT, XRCC1, and XPD and cancer risk in genetic epidemiology. Methods Mol Biol. 2009;471:305–33. doi: 10.1007/978-1-59745-416-2_16. PubMed PMID: 19109787.

7. Patrono C, Sterpone S, Testa A, Cozzi R. Polymorphisms in base excision repair genes: Breast cancer risk and individual radiosensitivity. World J Clin Oncol. 2014;5(5):874–82. doi: 10.5306/wjco.v5.i5.874. PubMed PMID: 25493225; PubMed Central PMCID: PMCPMC4259949.

8. Caldecott KW. DNA single-strand break repair. Experimental cell research. 2014;329(1):2–8. doi: 10.1016/j.yexcr.2014.08.027. PubMed PMID: 25176342.

9. Wilson SH, Kunkel TA. Passing the baton in base excision repair. Nat Struct Biol. 2000;7(3):176–8. doi: 10.1038/73260. PubMed PMID: 10700268.

10. Audebert M, Salles B, Calsou P. Involvement of poly(ADP-ribose) polymerase-1 and XRCC1/DNA ligase III in an alternative route for DNA double-strand breaks rejoining. J Biol Chem. 2004;279(53):55117–26. doi: 10.1074/jbc.M404524200. PubMed PMID: 15498778.

11. Frit P, Barboule N, Yuan Y, Gomez D, Calsou P. Alternative end-joining pathway(s): bricolage at DNA breaks. DNA Repair (Amst). 2014;17:81–97. doi: 10.1016/j.dnarep.2014.02.007. PubMed PMID: 24613763.

12. Mladenov E, Iliakis G. Induction and repair of DNA double strand breaks: the increasing spectrum of non-homologous end joining pathways. Mutat Res. 2011;711(1-2):61–72. doi: 10.1016/j.mrfmmm.2011.02.005. PubMed PMID: 21329706.

13. Moser J, Kool H, Giakzidis I, Caldecott K, Mullenders LH, Fousteri MI. Sealing of chromosomal DNA nicks during nucleotide excision repair requires XRCC1 and DNA ligase III alpha in a cell-cycle-specific manner. Molecular cell. 2007;27(2):311–23. doi: 10.1016/j.molcel.2007.06.014. PubMed PMID: 17643379.

14. Narciso L, Fortini P, Pajalunga D, Franchitto A, Liu P, Degan P, et al. Terminally differentiated muscle cells are defective in base excision DNA repair and hypersensitive to oxygen injury. Proc Natl Acad Sci U S A. 2007;104(43):17010–5. doi: 10.1073/pnas.0701743104. PubMed PMID: 17940040; PubMed Central PMCID: PMCPMC2040456.

15. Sykora P, Yang JL, Ferrarelli LK, Tian J, Tadokoro T, Kulkarni A, et al. Modulation of DNA base excision repair during neuronal differentiation. Neurobiol Aging. 2013;34(7):1717–27. doi: 10.1016/j.neurobiolaging.2012.12.016. PubMed PMID: 23375654; PubMed Central PMCID: PMCPMC5576894.

16. Sultana R, Abdel-Fatah T, Abbotts R, Hawkes C, Albarakati N, Seedhouse C, et al. Targeting XRCC1 deficiency in breast cancer for personalized therapy. Cancer Res. 2013;73(5):1621–34. doi: 10.1158/0008-5472.CAN-12-2929. PubMed PMID: 23253910.

17. De Summa S, Pinto R, Pilato B, Sambiasi D, Porcelli L, Guida G, et al. Expression of base excision repair key factors and miR17 in familial and sporadic breast cancer. Cell Death Dis. 2014;5:e1076. doi: 10.1038/cddis.2014.30. PubMed PMID: 24556691; PubMed Central PMCID: PMCPMC3944247.

18. Ali R, Al-Kawaz A, Toss MS, Green AR, Miligy IM, Mesquita KA, et al. Targeting PARP1 in XRCC1-Deficient Sporadic Invasive Breast Cancer or Preinvasive Ductal Carcinoma In Situ Induces Synthetic Lethality and Chemoprevention. Cancer Res. 2018;78(24):6818–27. doi: 10.1158/0008-5472.CAN-18-0633. PubMed PMID: 30297533.

19. Albarakati N, Abdel-Fatah TM, Doherty R, Russell R, Agarwal D, Moseley P, et al. Targeting BRCA1-BER deficient breast cancer by ATM or DNA-PKcs blockade either alone or in combination with cisplatin for personalized therapy. Mol Oncol. 2015;9(1):204–17. doi: 10.1016/j.molonc.2014.08.001. PubMed PMID: 25205036; PubMed Central PMCID: PMCPMC5528668.

20. Horton JK, Wilson SH. Strategic Combination of DNA-Damaging Agent and PARP Inhibitor Results in Enhanced Cytotoxicity. Front Oncol. 2013;3:257. doi: 10.3389/fonc.2013.00257. PubMed PMID: 24137565; PubMed Central PMCID: PMCPMC3786324.

21. Horton JK, Stefanick DF, Prasad R, Gassman NR, Kedar PS, Wilson SH. Base excision repair defects invoke hypersensitivity to PARP inhibition. Mol Cancer Res. 2014;12(8):1128–39. doi: 10.1158/1541-7786.MCR-13-0502. PubMed PMID: 24770870; PubMed Central PMCID: PMCPMC4135006.

22. Vogelstein B, Kinzler KW. Cancer genes and the pathways they control. Nat Med. 2004;10(8):789–99. doi: 10.1038/nm1087. PubMed PMID: 15286780.

23. Chaim IA, Nagel ZD, Jordan JJ, Mazzucato P, Ngo LP, Samson LD. In vivo measurements of interindividual differences in DNA glycosylases and APE1 activities. Proc Natl Acad Sci U S A. 2017;114(48):E10379–E88. doi: 10.1073/pnas.1712032114. PubMed PMID: 29122935; PubMed Central PMCID: PMCPMC5715766.

24. Nagel ZD, Margulies CM, Chaim IA, McRee SK, Mazzucato P, Ahmad A, et al. Multiplexed DNA repair assays for multiple lesions and multiple doses via transcription inhibition and transcriptional mutagenesis. Proc Natl Acad Sci U S A. 2014;111(18):E1823–32. doi: 10.1073/pnas.1401182111. PubMed PMID: 24757057; PubMed Central PMCID: PMCPMC4020053.

25. Sonavane M, Sykora P, Andrews JF, Sobol RW, Gassman NR. Camptothecin Efficacy to Poison Top1 Is Altered by Bisphenol A in Mouse Embryonic Fibroblasts. Chem Res Toxicol. 2018;31(6):510–9. doi: 10.1021/acs.chemrestox.8b00050. PubMed PMID: 29799191.

26. Kirby TW, Gassman NR, Smith CE, Pedersen LC, Gabel SA, Sobhany M, et al. Nuclear Localization of the DNA Repair Scaffold XRCC1: Uncovering the Functional Role of a Bipartite NLS. Sci Rep. 2015;5:13405. doi: 10.1038/srep13405. PubMed PMID: 26304019; PubMed Central PMCID: PMCPMC4548243.

27. Kirby TW, Gassman NR, Smith CE, Zhao ML, Horton JK, Wilson SH, et al. DNA polymerase beta contains a functional nuclear localization signal at its N-terminus. Nucleic Acids Res. 2017;45(4):1958–70. doi: 10.1093/nar/gkw1257. PubMed PMID: 27956495; PubMed Central PMCID: PMCPMC5389473.

28. Gassman NR, Wilson SH. Micro-irradiation tools to visualize base excision repair and single-strand break repair. DNA Repair (Amst). 2015;31:52–63. doi: 10.1016/j.dnarep.2015.05.001. PubMed PMID: 25996408; PubMed Central PMCID: PMCPMC4458156.

29. Holton NW, Andrews JF, Gassman NR. Application of Laser Micro-irradiation for Examination of Single and Double Strand Break Repair in Mammalian Cells. J Vis Exp. 2017;(127). doi: 10.3791/56265. PubMed PMID: 28930988; PubMed Central PMCID: PMCPMC5752190.

30. Holton NW, Ebenstein Y, Gassman NR. Broad spectrum detection of DNA damage by Repair Assisted Damage Detection (RADD). DNA Repair (Amst). 2018;66-67:42–9. doi: 10.1016/j.dnarep.2018.04.007. PubMed PMID: 29723708.

31. Chandrashekar DS, Bashel B, Balasubramanya SAH, Creighton CJ, Ponce-Rodriguez I, Chakravarthi B, et al. UALCAN: A Portal for Facilitating Tumor Subgroup Gene Expression and Survival Analyses. Neoplasia. 2017;19(8):649–58. doi: 10.1016/j.neo.2017.05.002. PubMed PMID: 28732212; PubMed Central PMCID: PMCPMC5516091.

32. Hastak K, Alli E, Ford JM. Synergistic chemosensitivity of triple-negative breast cancer cell lines to poly(ADP-Ribose) polymerase inhibition, gemcitabine, and cisplatin. Cancer Res. 2010;70(20):7970–80. doi: 10.1158/0008-5472.CAN-09-4521. PubMed PMID: 20798217; PubMed Central PMCID: PMCPMC2955854.

33. Chavez KJ, Garimella SV, Lipkowitz S. Triple negative breast cancer cell lines: one tool in the search for better treatment of triple negative breast cancer. Breast Dis. 2010;32(1-2):35–48. doi: 10.3233/BD-2010-0307. PubMed PMID: 21778573; PubMed Central PMCID: PMCPMC3532890.

34. Boichuk S, Galembikova A, Sitenkov A, Khusnutdinov R, Dunaev P, Valeeva E, et al. Establishment and characterization of a triple negative basal-like breast cancer cell line with multi-drug resistance. Oncol Lett. 2017;14(4):5039–45. doi: 10.3892/ol.2017.6795. PubMed PMID: 29085518; PubMed Central PMCID: PMCPMC5649570.

35. Hui L, Zheng Y, Yan Y, Bargonetti J, Foster DA. Mutant p53 in MDA-MB-231 breast cancer cells is stabilized by elevated phospholipase D activity and contributes to survival signals generated by phospholipase D. Oncogene. 2006;25(55):7305–10. doi: 10.1038/sj.onc.1209735. PubMed PMID: 16785993.

36. Lim LY, Vidnovic N, Ellisen LW, Leong CO. Mutant p53 mediates survival of breast cancer cells. Br J Cancer. 2009;101(9):1606–12. doi: 10.1038/sj.bjc.6605335. PubMed PMID: 19773755; PubMed Central PMCID: PMCPMC2778523.

37. Nigro JM, Baker SJ, Preisinger AC, Jessup JM, Hostetter R, Cleary K, et al. Mutations in the p53 gene occur in diverse human tumour types. Nature. 1989;342(6250):705–8. doi: 10.1038/342705a0. PubMed PMID: 2531845.

38. Hollestelle A, Nagel JH, Smid M, Lam S, Elstrodt F, Wasielewski M, et al. Distinct gene mutation profiles among luminal-type and basal-type breast cancer cell lines. Breast Cancer Res Treat. 2010;121(1):53–64. doi: 10.1007/s10549-009-0460-8. PubMed PMID: 19593635.

39. O’Connor PM, Jackman J, Bae I, Myers TG, Fan S, Mutoh M, et al. Characterization of the p53 tumor suppressor pathway in cell lines of the National Cancer Institute anticancer drug screen and correlations with the growth-inhibitory potency of 123 anticancer agents. Cancer Res. 1997;57(19):4285–300. PubMed PMID: 9331090.

40. Gazdar AF, Kurvari V, Virmani A, Gollahon L, Sakaguchi M, Westerfield M, et al. Characterization of paired tumor and non-tumor cell lines established from patients with breast cancer. Int J Cancer. 1998;78(6):766–74. PubMed PMID: 9833771.

41. Fang Q, Inanc B, Schamus S, Wang XH, Wei L, Brown AR, et al. HSP90 regulates DNA repair via the interaction between XRCC1 and DNA polymerase beta. Nat Commun. 2014;5:5513. doi: 10.1038/ncomms6513. PubMed PMID: 25423885; PubMed Central PMCID: PMCPMC4246423.

42. Horton JK, Stefanick DF, Gassman NR, Williams JG, Gabel SA, Cuneo MJ, et al. Preventing oxidation of cellular XRCC1 affects PARP-mediated DNA damage responses. DNA Repair (Amst). 2013;12(9):774–85. doi: 10.1016/j.dnarep.2013.06.004. PubMed PMID: 23871146; PubMed Central PMCID: PMCPMC3924596.

43. Parsons JL, Tait PS, Finch D, Dianova, II, Allinson SL, Dianov GL. CHIP-mediated degradation and DNA damage-dependent stabilization regulate base excision repair proteins. Mol Cell. 2008;29(4):477–87. doi: 10.1016/j.molcel.2007.12.027. PubMed PMID: 18313385.

44. Prasad R, Caglayan M, Dai DP, Nadalutti CA, Zhao ML, Gassman NR, et al. DNA polymerase beta: A missing link of the base excision repair machinery in mammalian mitochondria. DNA Repair (Amst). 2017;60:77–88. doi: 10.1016/j.dnarep.2017.10.011. PubMed PMID: 29100041.

45. Gassman NR, Stefanick DF, Kedar PS, Horton JK, Wilson SH. Hyperactivation of PARP triggers nonhomologous end-joining in repair-deficient mouse fibroblasts. PloS one. 2012;7(11):e49301. doi: 10.1371/journal.pone.0049301. PubMed PMID: 23145148; PubMed Central PMCID: PMCPMC3492265.

46. Slyskova J, Sabatella M, Ribeiro-Silva C, Stok C, Theil AF, Vermeulen W, et al. Base and nucleotide excision repair facilitate resolution of platinum drugs-induced transcription blockage. Nucleic Acids Res. 2018;46(18):9537–49. doi: 10.1093/nar/gky764. PubMed PMID: 30137419; PubMed Central PMCID: PMCPMC6182164.

47. Sawant A, Floyd AM, Dangeti M, Lei W, Sobol RW, Patrick SM. Differential role of base excision repair proteins in mediating cisplatin cytotoxicity. DNA Repair (Amst). 2017;51:46–59. doi: 10.1016/j.dnarep.2017.01.002. PubMed PMID: 28110804; PubMed Central PMCID: PMCPMC5328804.

48. Prasad R, Horton JK, Dai DP, Wilson SH. Repair pathway for PARP-1 DNA-protein crosslinks. DNA Repair (Amst). 2018. doi: 10.1016/j.dnarep.2018.11.004. PubMed PMID: 30466837.

49. Kang YJ, Yan CT. Regulation of DNA repair in the absence of classical non-homologous end joining. DNA Repair (Amst). 2018;68:34–40. doi: 10.1016/j.dnarep.2018.06.001. PubMed PMID: 29929045.

50. Lehmann BD, Bauer JA, Chen X, Sanders ME, Chakravarthy AB, Shyr Y, et al. Identification of human triple-negative breast cancer subtypes and preclinical models for selection of targeted therapies. J Clin Invest. 2011;121(7):2750–67. doi: 10.1172/JCI45014. PubMed PMID: 21633166; PubMed Central PMCID: PMCPMC3127435.

51. Yang W, Soares J, Greninger P, Edelman EJ, Lightfoot H, Forbes S, et al. Genomics of Drug Sensitivity in Cancer (GDSC): a resource for therapeutic biomarker discovery in cancer cells. Nucleic Acids Res. 2013;41(Database issue):D955–61. doi: 10.1093/nar/gks1111. PubMed PMID: 23180760; PubMed Central PMCID: PMCPMC3531057.

52. Horton JK, Watson M, Stefanick DF, Shaughnessy DT, Taylor JA, Wilson SH. XRCC1 and DNA polymerase beta in cellular protection against cytotoxic DNA single-strand breaks. Cell Res. 2008;18(1):48–63. doi: 10.1038/cr.2008.7. PubMed PMID: 18166976; PubMed Central PMCID: PMCPMC2366203.

53. Pommier Y, O’Connor MJ, de Bono J. Laying a trap to kill cancer cells: PARP inhibitors and their mechanisms of action. Sci Transl Med. 2016;8(362):362ps17. doi: 10.1126/scitranslmed.aaf9246. PubMed PMID: 27797957.

54. Gassman NR, Coskun E, Stefanick DF, Horton JK, Jaruga P, Dizdaroglu M, et al. Bisphenol a promotes cell survival following oxidative DNA damage in mouse fibroblasts. PLoS One. 2015;10(2):e0118819. doi: 10.1371/journal.pone.0118819. PubMed PMID: 25693136; PubMed Central PMCID: PMCPMC4334494.

55. Whitaker A, Schaich M, Smith M, Flynn T, Freudenthal B. Base excision repair of oxidative DNA damage: from mechanism to disease. Front Biosci (Landmark Ed). 2017;22:1493–522. PubMed Central PMCID: PMCPMC5567671.

56. Uhlen M, Zhang C, Lee S, Sjostedt E, Fagerberg L, Bidkhori G, et al. A pathology atlas of the human cancer transcriptome. Science. 2017;357(6352). doi: 10.1126/science.aan2507. PubMed PMID: 28818916.

57. Masson M, Niedergang C, Schreiber V, Muller S, Menissier-de Murcia J, de Murcia G. XRCC1 is specifically associated with poly(ADP-ribose) polymerase and negatively regulates its activity following DNA damage. Mol Cell Biol. 1998;18(6):3563–71. PubMed PMID: 9584196; PubMed Central PMCID: PMCPMC108937.

58. Kiriyama T, Hirano M, Asai H, Ikeda M, Furiya Y, Ueno S. Restoration of nuclear-import failure caused by triple A syndrome and oxidative stress. Biochem Biophys Res Commun. 2008;374(4):631–4. doi: 10.1016/j.bbrc.2008.07.088. PubMed PMID: 18662670.

59. Lakshmipathy U, Campbell C. Mitochondrial DNA ligase III function is independent of Xrcc1. Nucleic Acids Res. 2000;28(20):3880–6.

60. Gao Y, Katyal S, Lee Y, Zhao J, Rehg JE, Russell HR, et al. DNA ligase III is critical for mtDNA integrity but not Xrcc1-mediated nuclear DNA repair. Nature. 2011;471(7337):240–4. doi: 10.1038/nature09773. PubMed PMID: 21390131; PubMed Central PMCID: PMCPMC3079429.

61. Takanami T, Nakamura J, Kubota Y, Horiuchi S. The Arg280His polymorphism in X-ray repair cross-complementing gene 1 impairs DNA repair ability. Mutat Res. 2005;582(1-2):135–45. doi: 10.1016/j.mrgentox.2005.01.007. PubMed PMID: 15781218.

62. Braithwaite EK, Kedar PS, Lan L, Polosina YY, Asagoshi K, Poltoratsky VP, et al. DNA polymerase lambda protects mouse fibroblasts against oxidative DNA damage and is recruited to sites of DNA damage/repair. J Biol Chem. 2005;280(36):31641–7. doi: 10.1074/jbc.C500256200. PubMed PMID: 16002405.

63. Braithwaite EK, Prasad R, Shock DD, Hou EW, Beard WA, Wilson SH. DNA polymerase lambda mediates a back-up base excision repair activity in extracts of mouse embryonic fibroblasts. J Biol Chem. 2005;280(18):18469–75. doi: 10.1074/jbc.M411864200. PubMed PMID: 15749700.

64. Polo LM, Xu Y, Hornyak P, Garces F, Zeng Z, Hailstone R, et al. Efficient Single-Strand Break Repair Requires Binding to Both Poly(ADP-Ribose) and DNA by the Central BRCT Domain of XRCC1. Cell Rep. 2019;26(3):573–81 e5. doi: 10.1016/j.celrep.2018.12.082. PubMed PMID: 30650352; PubMed Central PMCID: PMCPMC6334254.

65. Camirand A, Fadhil I, Luco AL, Ochietti B, Kremer RB. Enhancement of taxol, doxorubicin and zoledronate anti-proliferation action on triple-negative breast cancer cells by a PTHrP blocking monoclonal antibody. Am J Cancer Res. 2013;3(5):500–8. PubMed PMID: 24224127; PubMed Central PMCID: PMCPMC3816969.

