## Supplementary figures and images for "Defective base excision repair in the response to DNA damaging agents in triple negative breast cancer"

### Supplemental Figure 1

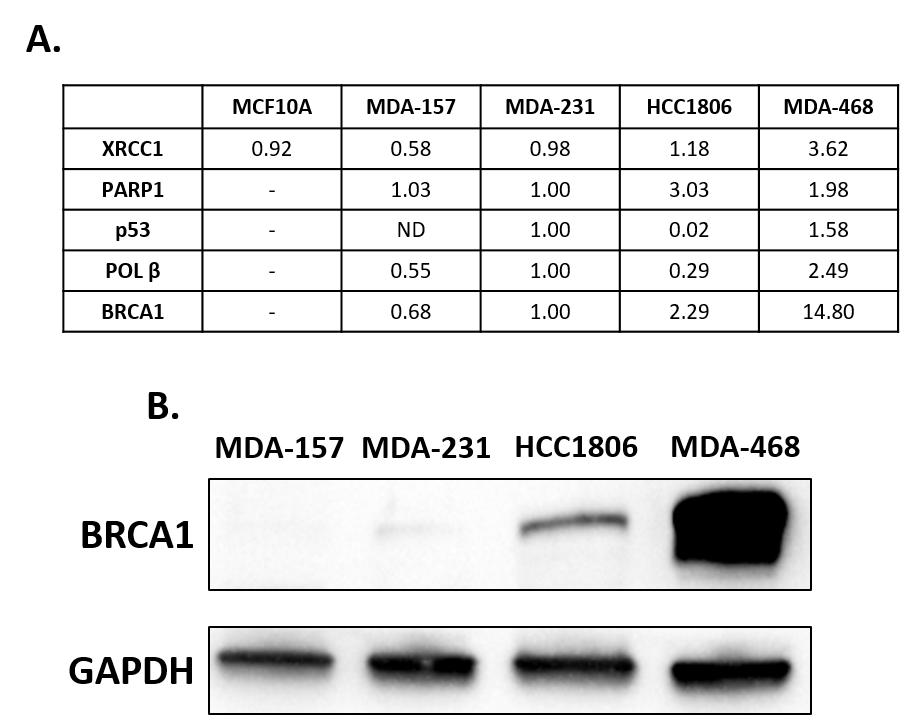

### Supplemental Figure 2

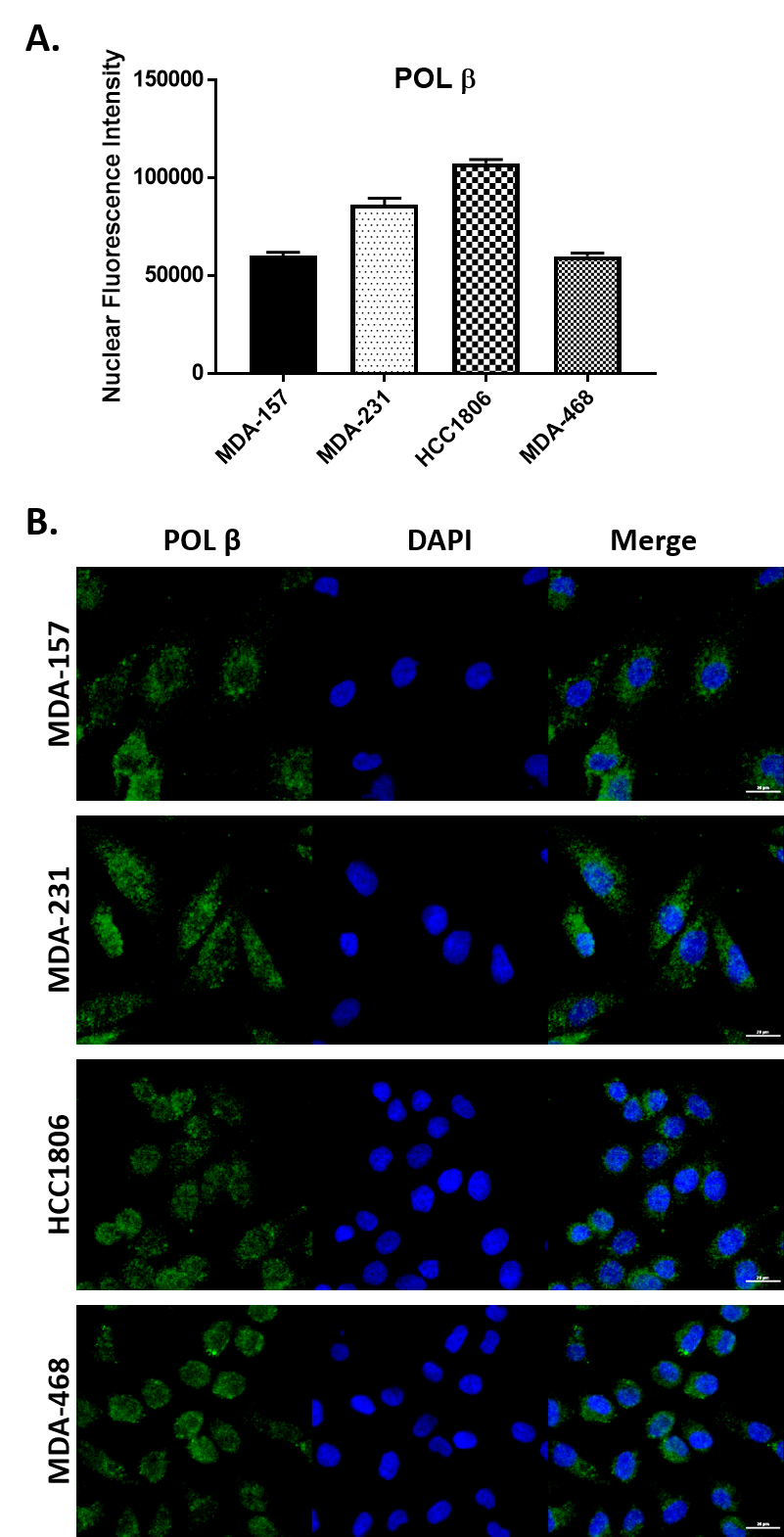

### Supplemental Figure 3

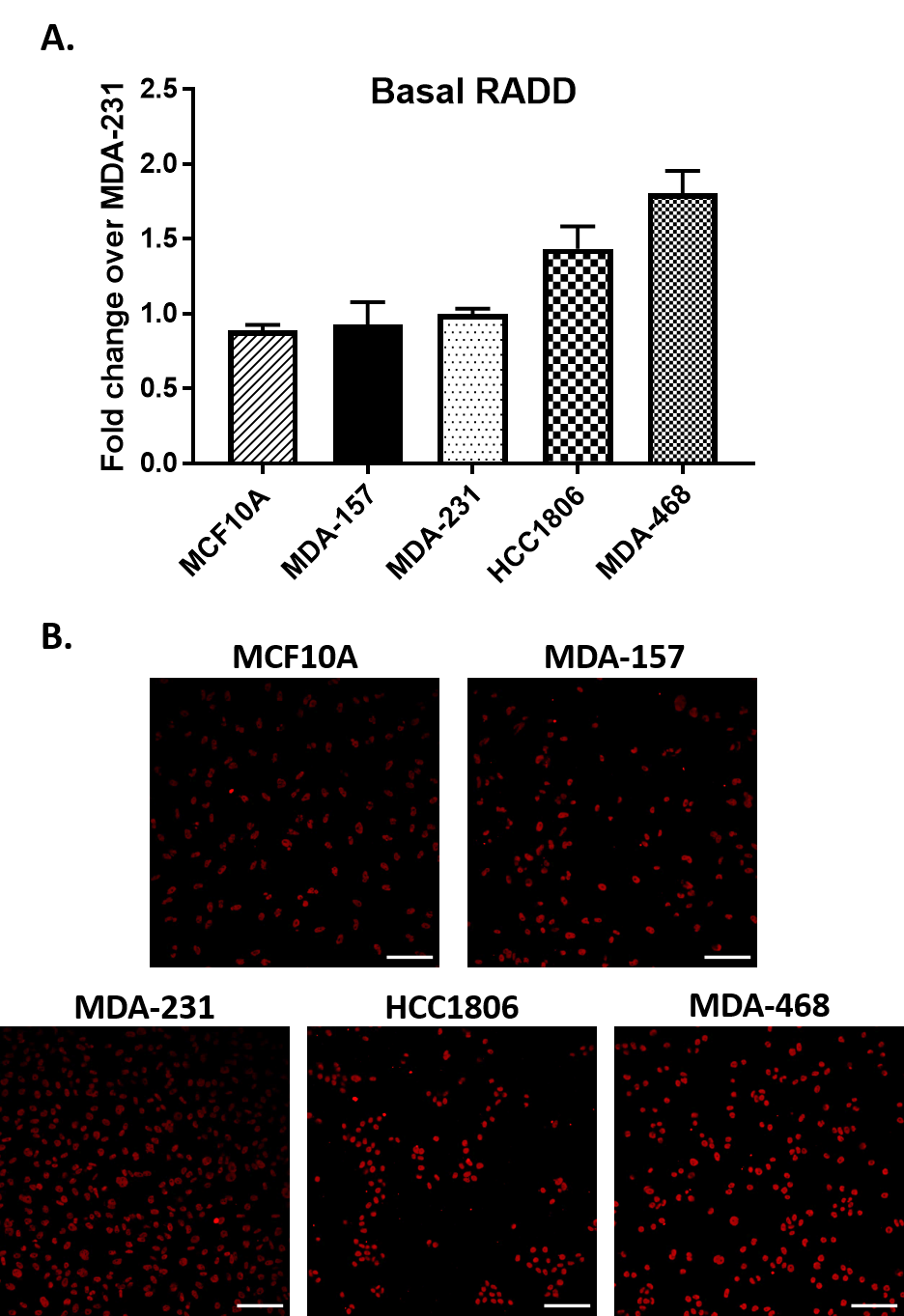

### Supplemental Figure 4

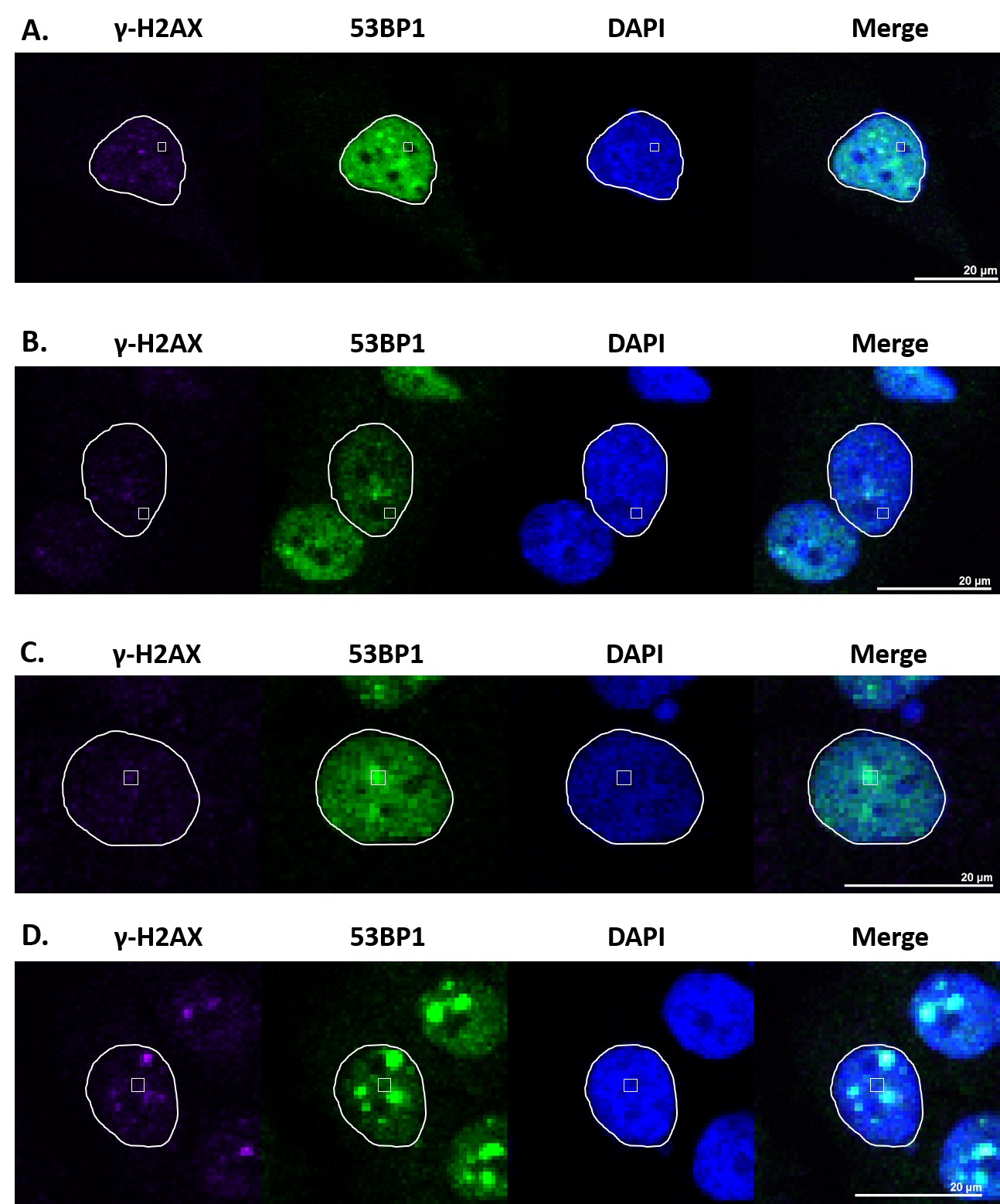

### Supplemental Figure 5

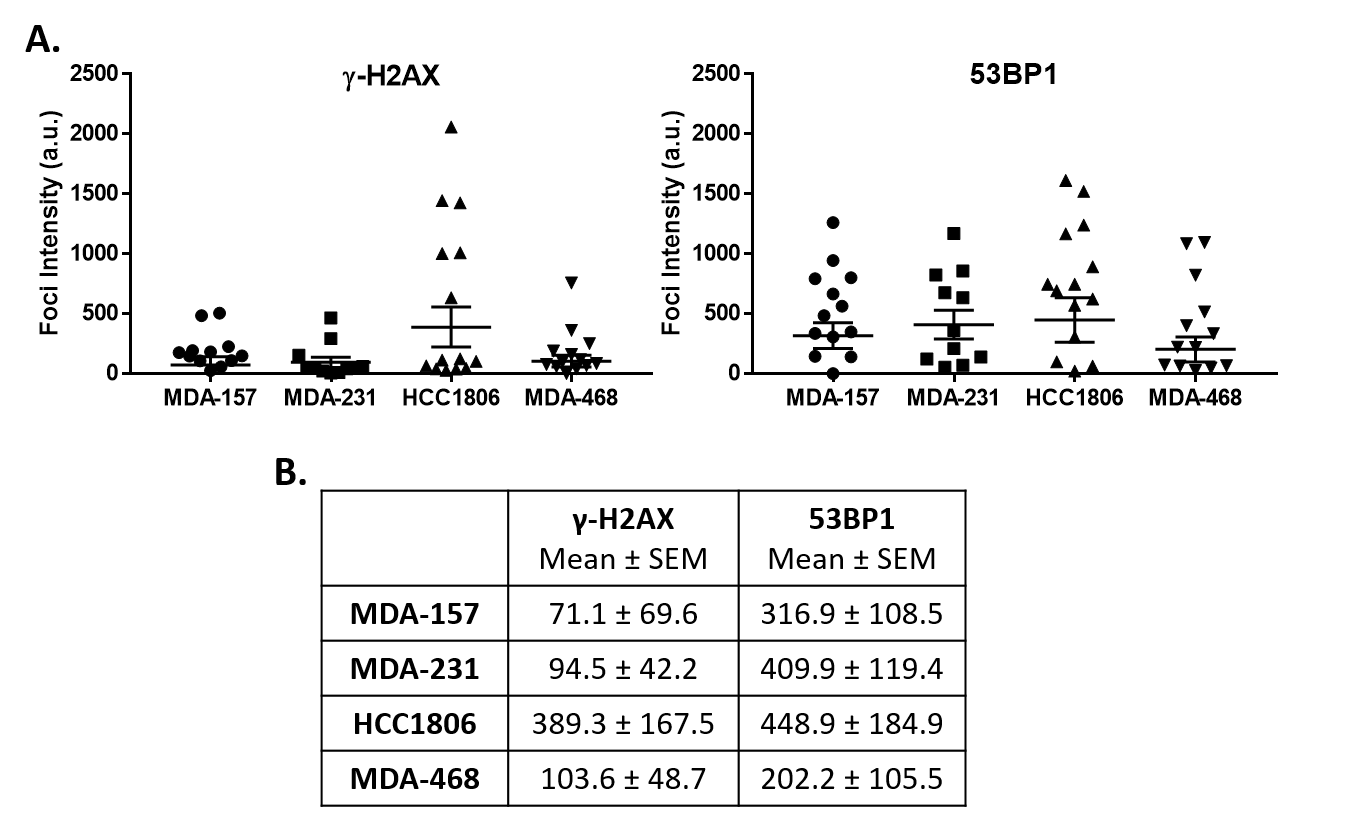

### Supplemental Figure 6

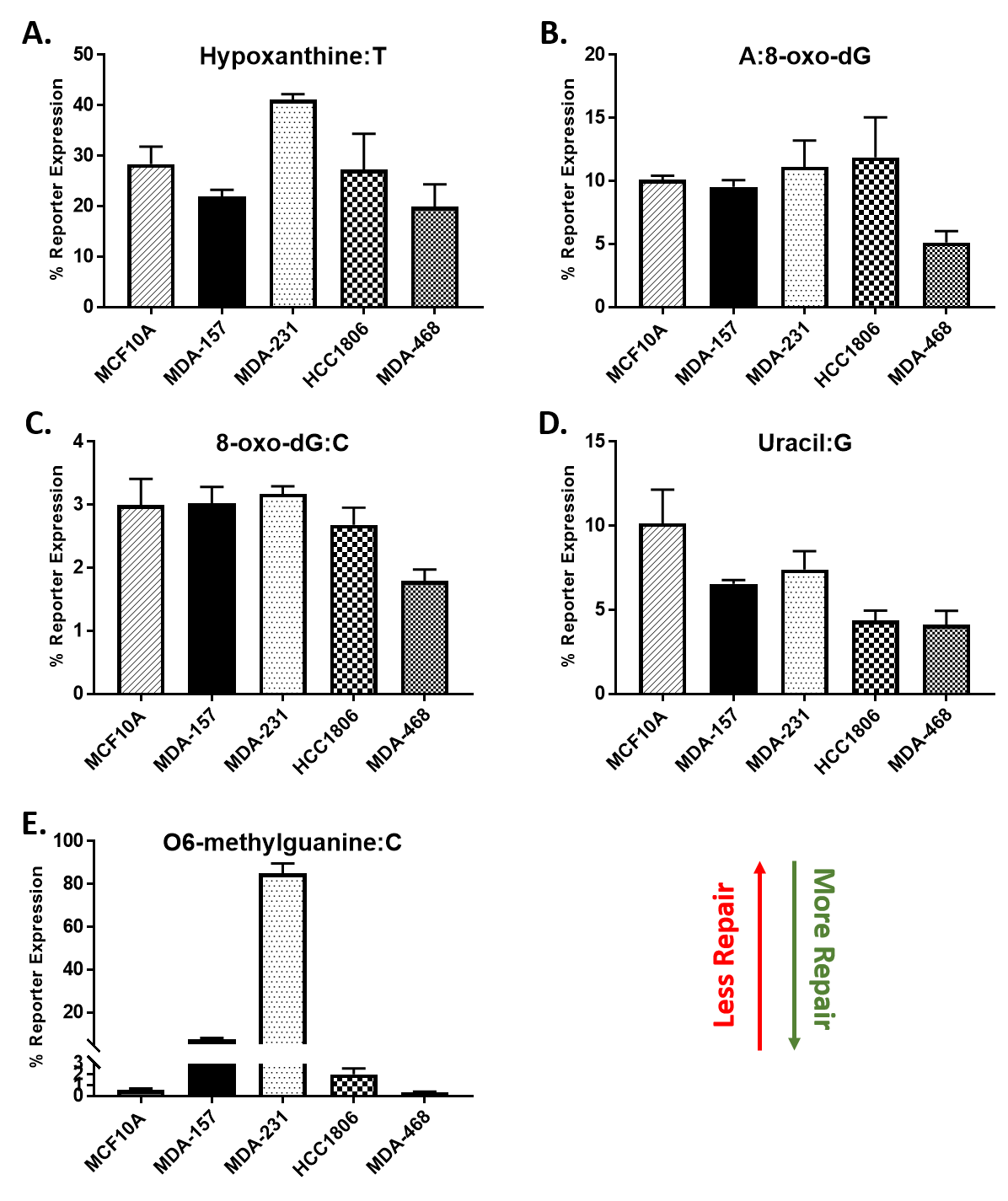
